# Curvature-based machine learning method for automated segmentation of dendritic spines

**DOI:** 10.64898/2025.12.04.692360

**Authors:** Abdel Kader A. Geraldo, Michael A. Chirillo, Kristen M. Harris, Thomas G. Fai

**Affiliations:** Department of Mathematics, Brandeis University, Waltham, MA, USA; Department of Biological Sciences, University of Rhode Island, Kingston, RI, USA; Department of Neuroscience, University of Texas at Austin, Austin, TX, USA; Department of Mathematics and Volen Center for Complex Systems, Brandeis University, Waltham, MA, USA

## Abstract

Recent advances in connectomics have been led by high-resolution reconstruction of large volumes of neural tissues using electron microscopy (EM), providing unprecedented insights into brain structure and function. Dendritic spines—dynamic protrusions on neuronal dendrites—play crucial roles in synaptic plasticity, influencing learning, memory, and various neurological disorders. However, current spine analysis methods often rely on manual annotation of subcellular features, limiting their ability to handle the complexity of spines in dense dendritic networks. This paper introduces a novel automated computational framework that integrates discrete differential geometry, machine learning, and 3D image processing to analyze dendritic spines in these intricate environments. By generating distributions of spine morphology from high resolution images including many thousands of spines, our approach captures subtle variations in spine shapes, offering a nuanced understanding of their roles in synaptic function. This framework is tested on multiple EM datasets, with the aim of enhancing our understanding of synaptic plasticity and its alterations in disease states. The proposed method is poised to accelerate neuroscience research by providing a scalable, objective, and comprehensive solution for spine analysis, uncovering insights into the role of spine geometry for neural function.

## 1. Introduction

Recent advances in electron microscopy and connectomics have made it possible to reconstruct astonishing large volumes of neural tissue at nanometer resolution (EM) [25, 29]. These breakthroughs have opened new opportunities for studying brain microstructures, particularly dendritic spines, dynamic protursion on dendrites where most excitatory synapses in the brain occur. The morphology of dendritic spines—encompassing shape, size, and density—is highly plastic and closely tied to fundamental neurological processes such as learning and memory [6, 19, 23, 46, 59].

Quantitative studies have shown that fine-scale geometric features of spines, including head volume, neck length, and curvature, are strongly associated with synaptic strength, developmental state, and functional specialization [37–39]. Changes in spine structure have also been linked to drug abuse, environmental influences, and a wide range of neurodevelopmental, neurodegenerative, and psychiatric disorders [16, 35, 57].

Accurate detection and analysis of dendritic spines in three-dimensional reconstructions is therefore essential for advancing our understanding of synaptic organization and plasticity. However, spine identification remains challenging due to their heterogeneous morphologies, dense clustering, and local curvature variations. Manual annotation is highly time-intensive and impractical for large-scale datasets, underscoring the need for automated and reliable computational methods. Several segmentation approaches have been proposed in recent years [4, 5, 41, 45, 47, 50, 52, 54, 56], yet challenges remain in achieving both accuracy and scalability.

In particular, many existing 3D CNN–based methods [2, 10, 34], operate directly on voxel representations, which require dense volumetric grids and therefore scale poorly to large EM datasets. This voxel dependence leads to substantial memory overhead, slow inference, and difficulty capturing fine geometric details such as thin spine necks. More broadly, large EM volumes contain millions of surface elements, creating significant computational constraints for voxel-level processing or repeated high-resolution inference. Together, these challenges underscore the need for automated approaches that incorporate geometric cues and remain robust across large, heterogeneous datasets.

To address these challenges, we introduce a method for automated spine segmentation that combines discrete differential geometry with deep learning. Our framework begins by preprocessing 3D reconstructions of dendritic segments to smooth the triangulated mesh, reduce noise, and enhance surface quality. From the resulting meshes, we extract geometric descriptors such as Gaussian and mean curvature, along with additional features including distances from the dendritic shaft skeleton and clustering-based descriptors. Together, these features provide a rich representation of both local and global morphology.

Building on this geometric foundation, we developed a series of deep neural network (DNN) architectures to evaluate how feature enrichment impacts segmentation performance. The baseline model, **DNN**_1_, relies primarily on curvature-based descriptors. **DNN**_2_ extends this by incorporating distance-to-skeleton features, improving the separation of spines from shafts. Finally, **DNN**_3_ integrates a set of enriched geometric and topological descriptors, enabling the network to capture subtle morphological variations and complex spine arrangements. This progression of models demonstrates how systematically incorporating new features enhances both training convergence and segmentation accuracy.

## 2. Methods and materials

In this section, we employ differential geometry to design a deep neural network architecture for the segmentation of dendritic spines. As a first step, we analyze how curvature can contribute to the characterization of dendritic morphology. Building on this analysis, we then develop and explore a deep neural network that leverages the geometric properties of dendrites to achieve accurate segmentation. For convenience, a summary table of the symbols and variables used throughout Sections 2 is provided in Supplemental Table S1.

### 2.1. Dendritic Morphology Analysis Using Discrete Differential Geometry

Segmentation of dendritic shafts and spines can be guided by their differential geometric properties, particularly Gaussian and mean curvature. These curvature measures capture local shape variations that distinguish the roughly cylindrical shaft from protruding spines. However, raw curvature values obtained directly from EM reconstructions are often noisy due to the sectioning process and mesh irregularities. To address this, we first smooth the dendritic triangulated surface mesh using discrete differential geometry techniques, then compute curvature values, and finally enhance these through image processing filters. This three-step process—smoothing, curvature computation, and enhancement–provides robust geometric descriptors that form the basis for accurate segmentation.

#### 2.1.1. Mean and Gaussian curvature

In this section, we motivate the use of Gaussian and mean curvature values as key variables for segmenting the dendritic shaft. In the study of dendritic morphology, the shaft is typically modeled as an approximately cylindrical structure from which spines protrude. The discrete mean **H** and Gaussian curvatures **K** computation details are provided in Section S5.

The Gaussian curvature provides insight into the surface’s shape at different points. At hyperbolic points, where the surface curves in opposite directions (saddle-like), the Gaussian curvature is negative. Conversely, at elliptical points, where the surface curves uniformly in the same direction (dome-like), the Gaussian curvature is positive. This contrasts with the cylindrical shaft, along which the Gaussian curvature is zero. Applying these principles to dendritic morphology, we observe that along the spine neck and at the intersection between the neck of a spine and the dendritic shaft, the surface exhibits a saddle shape, leading to negative Gaussian curvature. On the other hand, the spine head, with its more spherical or dome-like structure, exhibits positive Gaussian curvature.

The mean curvature further characterizes the surface. On a concave surface, the mean curvature is positive, while on a convex surface, it is negative. This distinction helps to identify regions of significant shape change. For instance, at the transition from the dendritic shaft to the spine neck, the surface is concave, leading to relatively positive mean curvature. In contrast, the spine head, which is a convex region, has relatively negative mean curvature. These curvature measures are most useful for segmentation when the triangular mesh is sufficiently smooth, as smoothness preserves the overall geometry of the dendritic branches while reducing discretization noise. Under these conditions, the curvature fields more accurately capture the cylindrical shape of the dendritic shaft and the geometric deviations introduced by spine necks and spine heads. As a result, the smoothed curvature values provide a more reliable representation of the underlying morphology, which is valuable in distinguishing the shaft from protruding spines.

#### 2.1.2. Smoothing the Triangulated Surface Mesh

To analyze the triangulated surface of the dendritic mesh, we first address the inherent roughness introduced by the EM sectioning process. We apply discrete differential–geometry techniques–specifically a discrete approximation of mean-curvature (Willmore) flow–to smooth the triangular surface mesh.

We define *k*_*bend*_ as the bending coefficient of the surface. The curvature energy over the surface *A* is

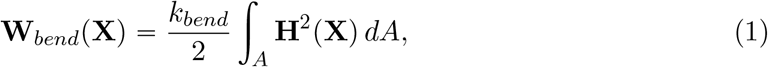

and the associated bending force is obtained as the negative gradient of this energy:

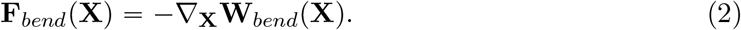

The surface is then evolved by a gradient-descent step,

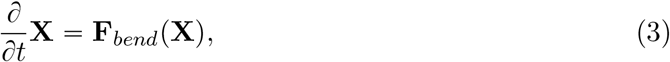

which produces a smoothed version of the mesh. In practice, this smoothing is implemented using standard discrete differential-geometry operators. Although the mesh consists of discrete triangles, smoothing is well-defined: each vertex exchanges curvature information only with the vertices in its 1-ring neighborhood (the set of triangles directly adjacent to it), as illustrated in Figure S1. Updating each vertex using curvature information from its neighbors provides a discrete analogue of smoothing on a continuous surface. This suppresses high-frequency numerical noise introduced by the discretization while preserving the underlying geometric structure of the dendritic mesh. Further numerical details, following the approach of [55], are provided in Section S5.

The results of performing this smoothing are illustrated in Figure 1, where we compute the Gaussian and mean curvature of a segment of spiny dendrite. The data was obtained from high-resolution 3D EM reconstructions of dendritic segments in the CA1 region of the hippocampus from P21 rats. Slices were subjected to *in vitro* theta-burst stimulation to induce LTP or to control stimulation, as described in [7–9]. Comparing the curvature before and after smoothing, we observe that the processed mesh exhibits a more easily interpretable profile of the dendritic surface, which can be exploited to improve the accuracy of spine-shaft segmentation.

**Figure 1.**
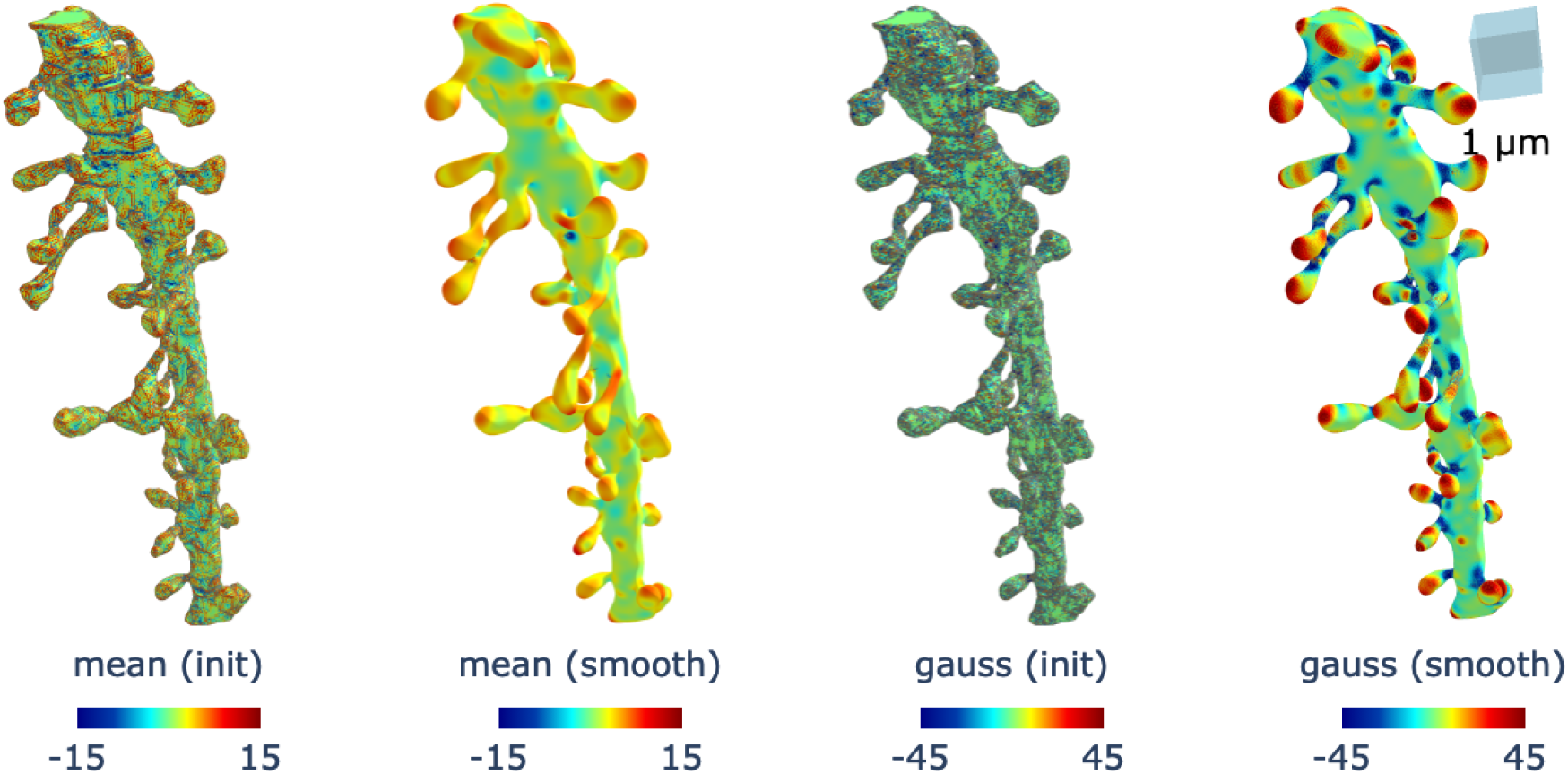
Effect of curvature smoothing on dendritic meshes. The plots labeled (init) and (smooth) show the initial dendritic curvature and the curvature after smoothing, respectively. The plots labeled mean and gauss correspond to the mean curvature and Gaussian curvature of a segment of dendrite. Smoothing enhances the curvature profile, producing a more easily interpretable pattern in the mesh that can be effectively leveraged to improve segmentation accuracy. For visualization, Gaussian curvature values are thresholded to remain within an absolute value of 45, while the mean curvature are constrained within an absolute value of 15, thereby highlighting the most relevant mesh faces. The triangular mesh is obtained from CA1 dendritic reconstructions [7–9].

#### 2.1.3. Enhancement of Curvature through Image Processing

Here, we provide an intuition for using image processing techniques to enhance the mean and Gaussian curvature values. The goal is to emphasize regions of the surface with significant curvature changes, which are more likely to correspond to dendritic shafts or spines, thereby facilitating segmentation. This forms the foundation for the machine learning methods that will be developed in the segmentation algorithm.

First, let us define the following sigmoidal function, which is often used in image processing to normalize and enhance contrast:

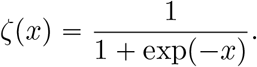

This transformation maps all real values into the interval (0, 1) . For large positive *x, ζ* (*x* ) →1, while for large negative *x, ζ* (*x* ) → 0. Around *x =* 0, the function has its steepest slope, which enhances small variations near zero and makes them more distinguishable.

Applying this transformation to the mean curvature **H** and the Gaussian curvature **K**, we obtain:

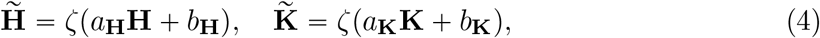

where *a*_**H**_, *b*_**H**_, *a*_**K**_, and *b*_**K**_ are empirically chosen parameters used to emphasize specific geometric features of the dendritic mesh. For example, in the neck region of a spine, we expect relatively high negative Gaussian curvature. By appropriately choosing *a*_**K**_ and *b*_**K**_, these negative values are pushed toward the lower end of the sigmoid, making them stand out more clearly during segmentation.

In contrast, along the cylindrical shaft, the Gaussian curvature is close to zero. Proper tuning ensures that values near zero are mapped consistently with the shaft, so that these regions are correctly identified. For mean curvature, which distinguishes concave and convex regions, the transformation highlights transitions: concave regions (positive mean curvature) are enhanced toward higher sigmoid values, while convex regions (negative mean curvature) are mapped toward lower values.

In practice, within this paper, the quantities 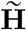 and 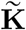 are not computed explicitly. Their formulation serves only to motivate the use of a DNN as an approximation function for the segmentation task, with a sigmoidal activation used in the output layer. The last two panels of Figure 3 show the spine and shaft probability outputs of the DNN that will be examined next. These results demonstrate a substantial improvement in segmentation quality compared to the manually tuned baseline. The overall processing pipeline—from mesh smoothing and geometric feature extraction to DNN-based prediction—is summarized in the flowchart shown in Figure S2.

### 2.2. Deep Neural Network Approach for Spine and Shaft Analysis

In the previous section, we provided an intuitive explanation of how segmentation can be enhanced using image processing techniques. We then introduced empirical filtering parameters that can improve segmentation quality. Instead of manually selecting these parameters and explicitly computing 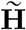 and 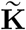, we can leverage Deep Neural Networks (DNNs) to learn an optimal segmentation approximation function based on **H** and **K**. At the same time, DNNs allow us to incorporate non-linearities that further enhance segmentation performance.

We analyze three different DNN architectures. The first network is designed to support the second by assisting in the extraction of the shaft skeleton, while the third network relies solely on external Python libraries for skeletonization. Machine learning methods have also been employed for dendrite segmentation of dendritic spines obtained from confocal reconstruction images using Convolution Neural Networks CNNs [52]. In this work, we adopt a simple DNN architecture inspired by the physics-informed neural network (PINN) model [22, 24, 30, 32, 33, 42–44], which has been widely applied to approximate ordinary and partial differential equations. These models leverage their well-known ability to serve as universal function approximators [22]. Given the geometrical characteristics of the dendritic triangular mesh, this approximation capability of DNNs constitutes a fundamental tool in our algorithm. Here, we present the architectures that produced the best results among the different trials.

#### 2.2.1. Deep Neural Network (**DNN**_1_) using the Gaussian and Mean Curvature

As for the sigmoidal function (4) introduced in the previous section, various model parameters require fine-tuning to enhance the shaft segmentation process. Rather than manually selecting these parameters and computing 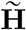 and 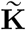, we train a deep neural network in a two-step procedure.

We first train a deep neural network, denoted as **DNN**_1_, whose architecture is illustrated in Figure 2. The input layer receives both the mean curvature **H** and the Gaussian curvature **K** values of the dendritic triangular mesh (after smoothing), together with their squared terms to capture higher-order variations. The input features are preprocessed such that the Gaussian curvature values and their squared terms are thresholded to remain within an absolute value of 45. Specifically, for **K** = ( **K**_1_, **K**_2_, …, **K**_*m*_ ), we enforce **K**_*i*_ ∈ [−45, 45 ]. We then compute 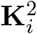 and threshold its value to the range [0, 45]. Similarly, the mean curvature values and their squared terms are constrained within an absolute value of 15, following the same procedure as in the Gaussian curvature case.

**Figure 2.**
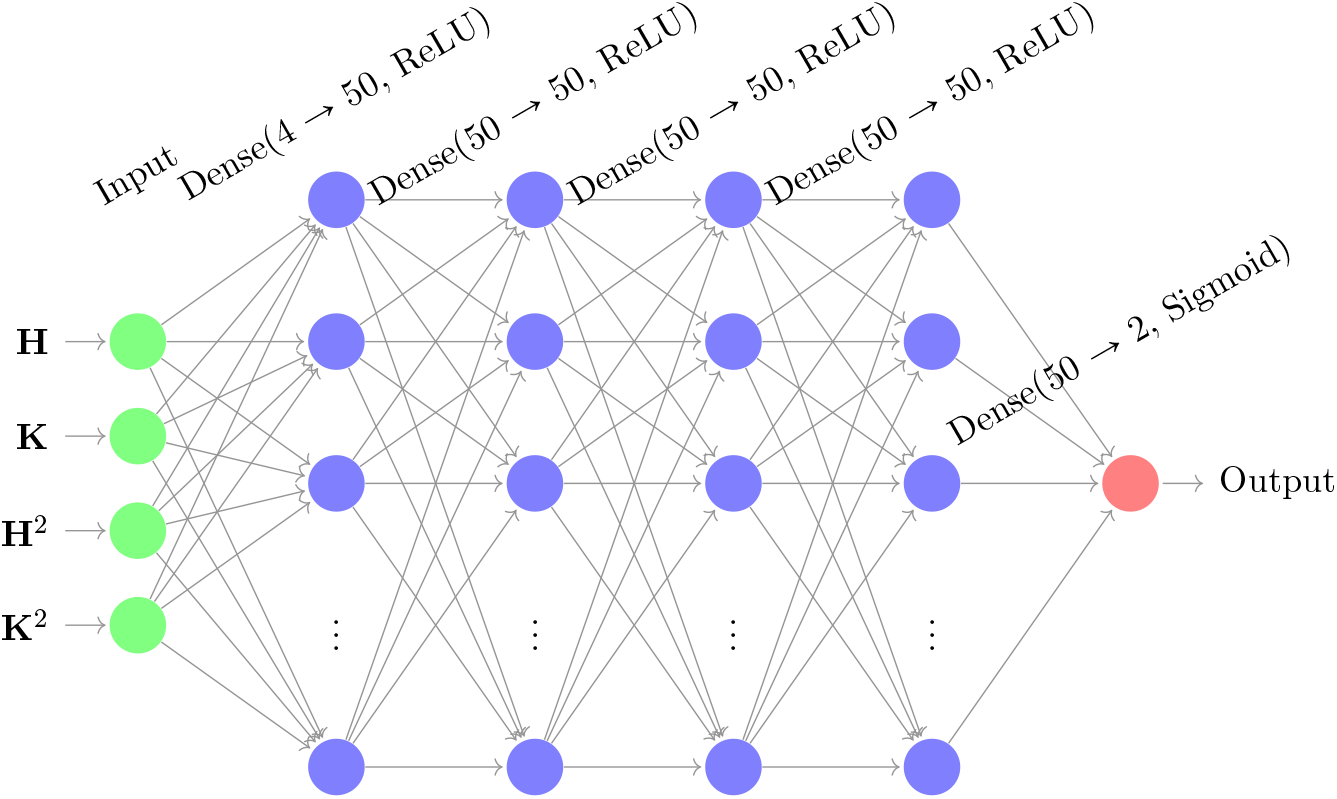
Deep Neural Network (**DNN**_1_) architecture used to segment the dendrite shaft. The input layer consists of the mean curvature **H** and the Gaussian curvature **K**, computed after smoothing, and their squares. The network includes four hidden layers, each with fifty neurons and ReLU activation functions. The output layer employs a sigmoidal activation function.

**Figure 3.**
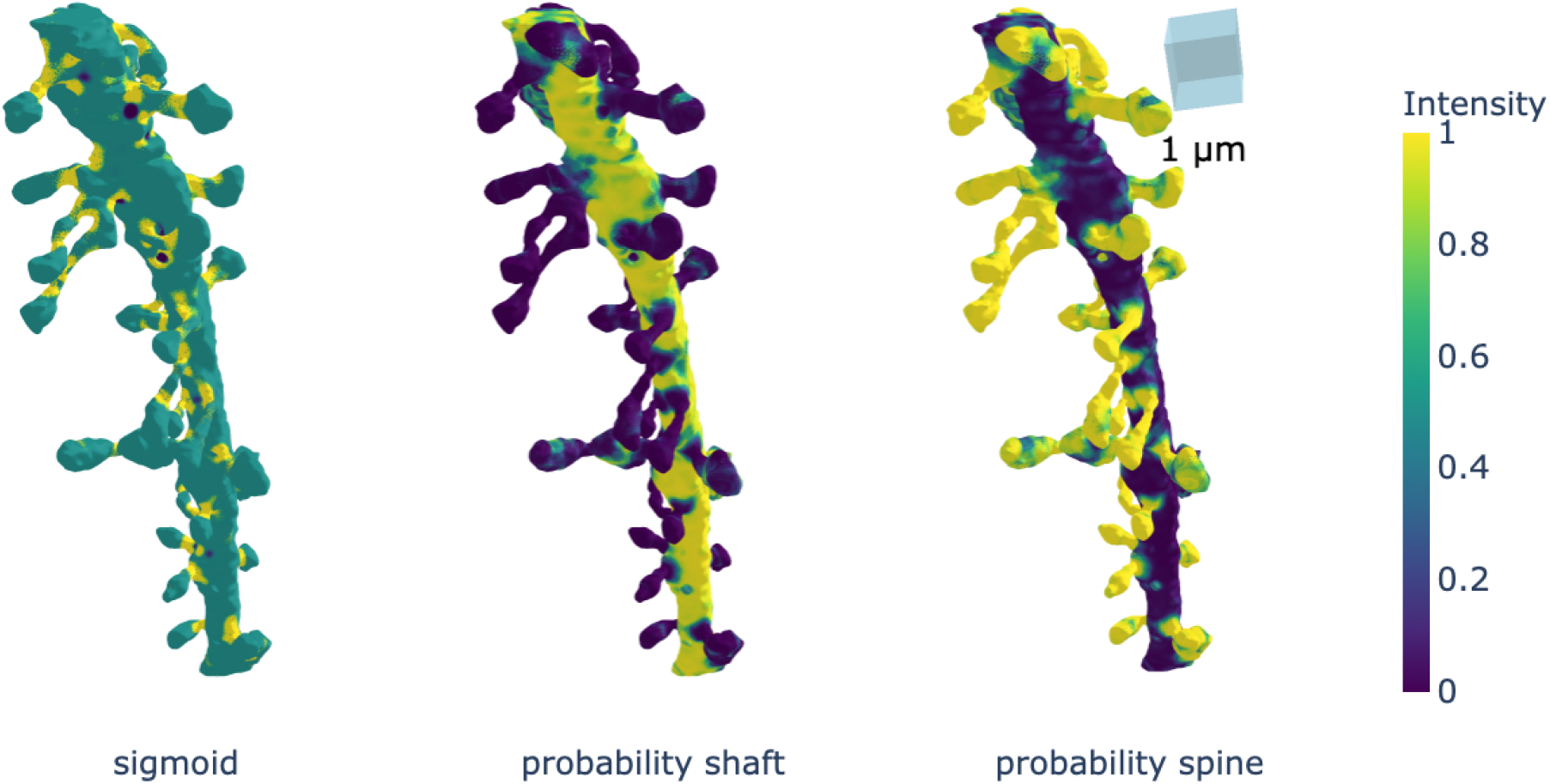
Visualization of geometric-based and learned probability representations used for dendritic spine segmentation. The panel labeled (sigmoid) shows the sigmoid of a linear combination of mean intensity and Gaussian curvature, which highlights the neck region and serves as a handcrafted baseline cue for segmentation. The panels labeled probability shaft and probability spine display the class-specific probability outputs of the network **DNN**_3_ for shaft and spine, respectively. Compared with the (sigmoid) representation, the learned probabilities provide a clearer and more discriminative separation of spine and shaft regions. The triangular mesh is obtained from CA1 dendritic reconstructions [7–9].

These thresholds are applied because curvature values can sometimes become very large, and such extreme values do not significantly improve prediction accuracy but can instead cause instability during training and inference. To further reduce instability, any NaN values, if present, are replaced with the average of the corresponding curvature values.

The network consists of four hidden layers, each containing fifty neurons with ReLU activation functions to introduce non-linearity [30]. The output layer employs a Sigmoid activation function, as described in Section 2.1.3, to classify vertices into dendritic shafts and spines.

The segmentation produced by **DNN**_1_ is not fully satisfactory, as regions within spines are sometimes misclassified as part of the shaft. This misclassification arises because certain spines regions exhibit relatively flat curvature, making them appear similar to shaft regions.

#### 2.2.2. Skeletonization

In this section, we describe the process used to obtain the skeletonization of dendritic meshes. is an image processing technique that reduces binary shapes to thin, single-pixel-wide lines while preserving their topological structure and connectivity [31, 60].

The first step in building the skeleton is to ensure that the mesh is *watertight*, meaning that it forms a completely closed surface with no gaps, holes, or disconnected edges. A watertight mesh guarantees a well-defined interior and exterior, which is essential for reliable geometric processing and for preserving the topological structure during skeletonization. To achieve this, we wrap the existing mesh using the algorithm described in Section S6.1. This procedure converts the surface into a uniformly sampled point cloud, estimates and orients normals, and then applies Poisson surface reconstruction to generate a closed, watertight representation.

In addition to Poisson reconstruction, we also evaluated the *alpha-wrap* method provided by `PyMeshLab` [11], which is based on the classical *α*-shape formulation [15]. Alpha wrapping produces well-behaved closures and can preserve fine geometric detail; however, in our experiments it was substantially slower than Poisson surface reconstruction for meshes of this size and resolution. For this reason, we adopted Poisson reconstruction as our primary wrapping method. For completeness, the alpha-wrap implementation is also included in Section S6.1.

Once a watertight mesh is obtained, we compute its skeleton using the `scikit-image` skeletonization package, which implements algorithms from [31,60]. Further implementation details are provided in Section S6. This simplified representation captures the essential branching geometry of dendrites for subsequent segmentation and morphological analysis.

#### 2.2.3. Deep Neural Network (**DNN**_2_) using Shaft Skeleton

We can enhance dendritic spine–shaft segmentation by incorporating the shaft skeleton. This involves using the distance between the shaft skeleton vertices and the dendritic branch as an additional input to the DNN. For this purpose, we use the shaft segmented with **DNN**_1_ together with the skeletonization procedure outlined in Section 2.2.2.

To begin, we consider the shaft-segmented meshes obtained from **DNN**_1_. Since this mesh results from the dendritic branch after spine regions have been segmented, it is no longer watertight. Therefore, we apply the procedure described earlier to make the mesh watertight. Once the shaft mesh skeletonization is completed, we compute the distance **D** between the shaft skeleton and the dendritic branch mesh vertices. Here, for each vertex **X**_*l*_, *l* ∈ { 1, 2, …, *n* } in the dendritic triangular mesh with *n* vertices, and { **V**_1_, **V**_2_, …, **V**_*v*_ } the set of shaft skeleton vertices, we compute the Euclidean norm:

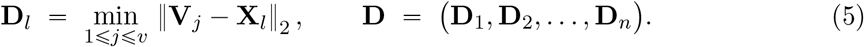

With this additional information, we develop an improved deep neural network, **DNN**_2_, whose architecture is shown in Figure S3. This model is similar to **DNN**_1_, except that it incorporates the new input feature. As in the earlier model, the input layer receives both the Gaussian and mean curvature values of the dendrite, along with the squared values of these curvatures as additional features. The network consists of four hidden layers, each containing fifty neurons with ReLU activation functions to introduce non-linearity. The output layer employs a Sigmoid activation function, as discussed in Section 2.1.3, to classify vertices into dendritic shafts or spines.

#### 2.2.4. Dendrite Spine-Shaft Segmentation Using Dendritic Branch Regions (**DNN**_3_)

To improve segmentation accuracy, we introduce a deep neural network (**DNN**_3_) that incorporates regional information from dendritic branch segments. These segments are derived from the dendrite skeleton, which serves as a structural reference for spatial organization. For this model, once the skeleton is extracted, we compute the shortest distance from each mesh vertex to its nearest skeleton point. These distances are then clustered using the K-means algorithm to partition the dendritic mesh into multiple regions. This regional segmentation enables us to distinguish between different parts of the dendrite shaft, spine neck, and spine head—regions that are critical for accurate classification.

As shown in Figure 4, to capture variations in dendritic morphology we apply K-means clustering with multiple values of *k*. We denote these segmentation features by **S**^*k*^, where *k* ∈ { 2, 3, …, 10 } corresponds to the number of clusters. In particular, the value of each feature **S**^*k*^ is given by the corresponding cluster label. Therefore this model incorporates 9 additional scalar features. In this study, we explore values of *k* ranging from 2 to 10 to provide a multi-scale representation of regional structure.

**Figure 4.**
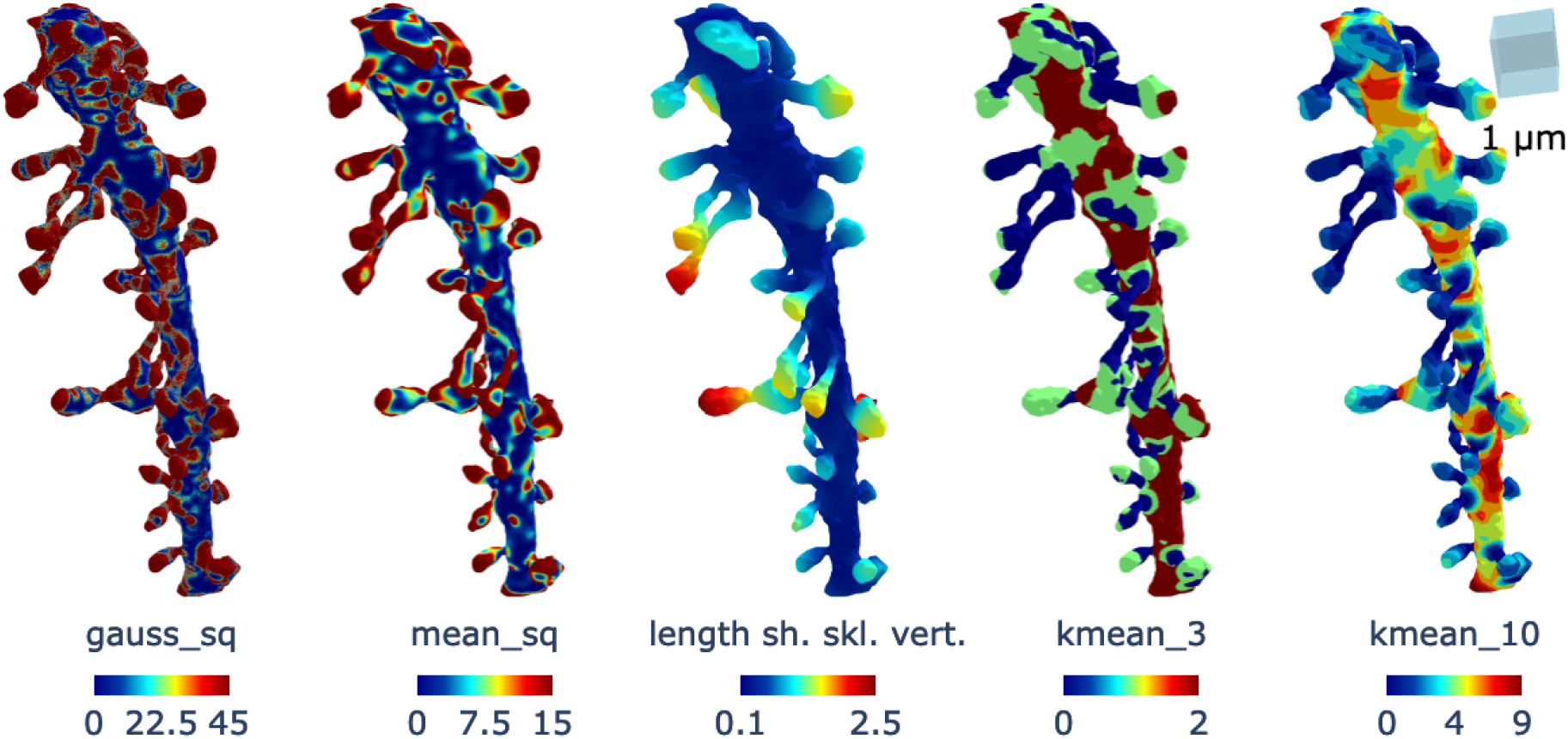
Additional features used in the DNN training. The plot labeled gauss_sq and mean_sq show respectively the square of the Gaussian and mean curvature of the dendrite. The plot labeled length sh. skl. vert. indicates the distance from the shaft skeleton to the dendritic mesh vertices, denoted by **D**. The plot labeled kmean_*k* illustrates the *k*-subdivision from K-means clustering, corresponding to the set **S**^*k*^. The triangular mesh is obtained from CA1 dendritic reconstructions [7–9].

In addition to these region-based segmentation features, we incorporate Gaussian curvature and its squared value as geometric descriptors. These measures are particularly effective for identifying neck regions, as discussed in earlier sections. We intentionally exclude mean curvature as we have observed that the mean curvature input increases the probability of misclassifying all dome-like regions as spines, leading to false positives. By omitting mean curvature and focusing on more discriminative features, **DNN**_3_ achieves improved segmentation performance compared to the baseline models.

Note that in practice, the K-means algorithm is not deterministic because its initial values depend on a random seed. To mitigate this and obtain consistent region assignments, we perform 50 independent clustering runs and select the labeling corresponding to the run with the lowest final K-means inertia. Since inertia measures the within-cluster compactness optimized by K-means, choosing the run with minimal inertia yields the most stable and coherent clustering among all runs. This procedure effectively filters out seed-dependent fluctuations and provides reproducible region assignments without requiring post hoc consensus voting.

We also considered using the silhouette score as an alternative selection criterion; however, computing silhouette values for all vertices is prohibitively expensive due to the large number of mesh vertices. For this reason, inertia-based selection offered a practical and computationally efficient solution.

### 2.3. Loss Function

We use the binary cross-entropy (BCE) to compute the loss function, and its derivation is presented below.

Let us consider a set of *m* dendritic meshes, denoted as 𝒟 = ( 𝒟_1_, 𝒟_2_, …, 𝒟_*m*_), where each mesh 𝒟_*i*_ consists of *n*_*i*_ vertices. For the training set, we assume that each 𝒟_*i*_ has a ground-truth classification matrix 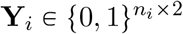, which represents the classification of vertices in 𝒟_*i*_ in one-hot encoding, such that:

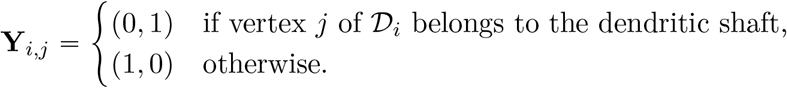

We define **Y** = ( **Y**_1_, **Y**_2_, …, **Y**_*m*_) as the set of all annotation matrices. Next, we define the set of mean curvature vectors for all dendritic meshes as **H** = (**H**_1_, **H**_2_, …, **H**_*m*_) and the set of Gaussian curvature vectors as **K** = (**K**_1_, **K**_2_, …, **K**_*m*_). For each mesh 𝒟_*j*_, *j* ∈ {1, 2, …, *m*}, the mean curvature vector is given by 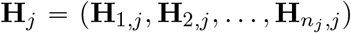, and the Gaussian curvature vector by 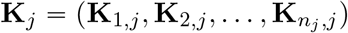, where **H**_*i,j*_ and **K**_*i,j*_ denote the mean and Gaussian curvatures of 𝒟_*j*_ at vertex *i*, for *i* ∈ {1, 2, …, *n*_*j*_}.

Additionally, we define the distance vector **D= (D**_1_, **D**_2_, …, **D**_*m*_ ), representing the distances between the central curve and the vertices, as well as additional structural descriptors **S**^*k*^, where *k* ∈ {2, 3, …, 10}. For each mesh 𝒟_*j*_, we write 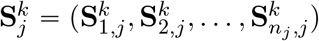.

We now define the feature vector for vertex *i* of mesh *j* as

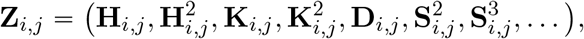

and the corresponding feature matrix for dendritic mesh 𝒟_*j*_ as

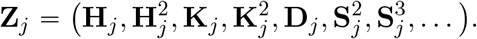

Our objective is to find a function

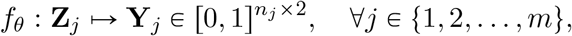

that best approximates the ground-truth classification by minimizing the following loss function:

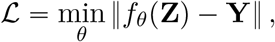

where ∥ · ∥ denotes an appropriate norm measuring the discrepancy between the predicted and actual classifications. In particular, we use a weighted sum of the binary cross-entropy (BCE), which encourages a more balanced optimization process. The binary cross-entropy formula is given by:

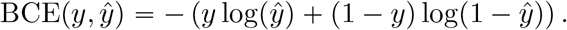

The final loss function is defined as:

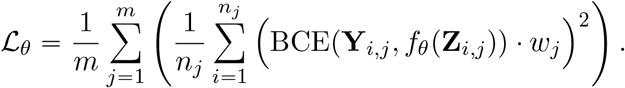

Here the weights *w*_*j*_ is empirically determined.

Note that the mean curvature **H** and Gaussian curvature **K** used during training are computed after smoothing the curvature fields on the triangular dendritic-branch meshes. The sole purpose of this smoothing is to obtain stable and interpretable curvature estimates that enhance the segmentation performance during training. As described in Section 2.1.2, smoothing the curvature improves the readability of the curvature profile and reduces noise-driven artifacts. Thus, the smoothing step in our pipeline serves only as a feature-enhancement operation, not as a geometric modification of the underlying dendrite. Importantly, all manual annotations were performed on the actual, unsmoothed dendritic branches, ensuring that the ground-truth labels correspond to the true biological geometry. During training, the network is therefore supervised to segment the real dendritic shaft and spine structures. After training, all dimensional measurements and quantitative analyses are performed on the original triangulated meshes, not on the smoothed versions.

### 2.4. Spine and Shaft Detection

After the training step of the algorithm, the next stage of spine–shaft segmentation involves applying the model for classification as well as performing additional post-processing steps to first isolate the shaft and then the spines. In this section, we analyze the processes required to obtain a reliable segmentation.

#### 2.4.1. Grouping Dendritic Mesh Parts into Connected Vertices

After training the DNN, the first step is to classify the vertices of a given test dendritic mesh into *shaft* and *spine* categories. The resulting classification can then be grouped into connected components of spine vertices and shaft vertices.

As a consequence of the sigmoidal activation function applied to the final layer, the output of the DNN is a probability matrix 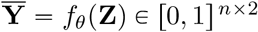, where each row corresponds to a vertex and contains the probabilities of that vertex being classified as a *spine*, 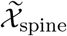 (first column greater than the second), or a *shaft*, 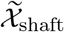 (second column greater than or equal to the first). Formally, we define:

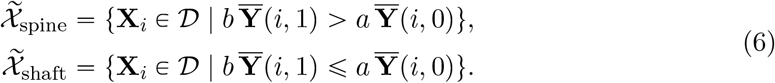

Here, *a* and *b* are user-defined weighting parameters that control the decision boundary. In our experiments, we fixed *b =* 0.81 and initially set *a =* 3. If the resulting segmentation produced too many shaft labels, we increased *a*; if too many spine labels were produced, we decreased *a*. This adjustment allowed us to fine-tune the spine–shaft separation based on the model’s output distribution.

Next, we describe how to group the predicted dendritic parts—classified as either *spine* or *shaft* —into subgroups of connected components. Let each vertex **X**_*i*_ have an associated set of neighboring vertices denoted by 𝒩_*i*_:

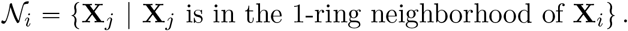

We define a *connected group* of vertices 𝒢_*i*_ ⊆ 𝒟 such that for any pair **X**_*p*_, **X**_*q*_ ∈ 𝒢_*i*_, there exists a path of vertices 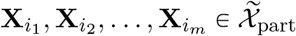 satisfying:

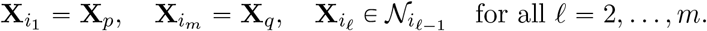

Here, 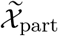 denotes the set of vertices classified as a given part type–either 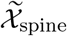 for spines or 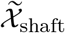 for shafts. The subset of *part-classified* vertices within this group is then:

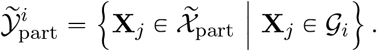

Finally, the complete set of part-classified vertices 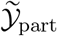 can be decomposed into disjoint set of connected vertices, where connectivity is defined by neighborhood overlap:

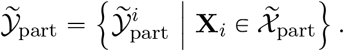

#### 2.4.2. Spine–Shaft Detection

Using the subgroups defined above of connected components, we now define the segmentation of the dendritic mesh into spines and shaft. Let 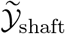 denote the set of connected components classified as *shaft*. In practice, some of these components may actually be parts of spines, so it is crucial to separate them from the true shaft set for accurate segmentation.

To achieve this, we identify the largest connected component in 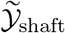 and designate it as the *entire shaft*. All remaining connected components in 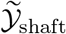 are then assumed to belong to spines.

The set of vertices belonging to the shaft is defined as:

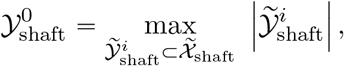

where | ·| denotes the cardinality of the set.

Once the entire shaft is defined, spine segmentation proceeds by removing the shaft vertices from the dendrite vertex set, reclassifying them as spine vertices and then reapplying the connected-component grouping process described in Section 2.4.1 to the remaining vertices.

The set of segmented spines is defined as:

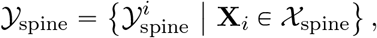

where each individual spine component is given by:

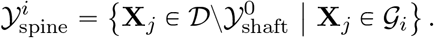

### 2.5. Dataset Description

This section describes the datasets of dendritic segments used for training and testing our algorithm. To mitigate overfitting, the training and testing sets were independent and originate from different animals. This separation ensures that the model learns generalizable features of dendritic structures rather than memorizing patterns specific to a single specimen. Table S3 provides a consolidated overview of the datasets used for training and testing, including their sources, annotation strategy, and selection criteria.

#### 2.5.1. Training dataset

The training dataset used as ground truth in this paper comprises six high-resolution 3D EM reconstructions of dendrites from the CA1 region of the hippocampus from rats at postnatal day 21 (P21), an immature developmental stage. Brain slices from these animals either underwent *in vitro* theta-burst stimulation to induce longterm potentiation (LTP) or received control stimulation, as described in [7–9].

For the training dataset, the meshes were manually annotated by experts, where spine and shaft segmentations were performed on all six dendritic branch reconstructions. Each spine was stored as an individual `.obj` file in watertight format. The mesh corresponding to the shaft of each dendritic segment was provided separately, along with the complete mesh of the dendritic segment. Because spine and shaft meshes were stored independently, we wrapped the spines and shaft with a tight mesh from the exterior in order to generate a unified dendritic mesh using the algorithm in Section S6.1. We then used the independent spine meshes to label the constituent parts of the new wrapped dendritic mesh. To achieve this, we employed a KD-Tree–based nearest-neighbor search to map spine vertices to the wrapped dendritic mesh. Specifically, for each set of spine vertices in the individual dataset `vertices`, we queried a KD-Tree built from the wrapped dendritic mesh vertices to identify all branch vertices within a radius threshold *r*_th_:

~~~
vertices_appr = list (set (np.concatenate (kdtree.query_radius(vertices, r_th_)))) .
~~~

Here, `query_radius` returns the indices of dendrite mesh vertices that lie within a distance *r*_th_ of each spine vertex. By aggregating these indices, we obtain the set of dendrite vertices corresponding to the annotated spine regions on the wrapped mesh. This mapping step allows us to merge the spine and shaft meshes into a single labeled dendritic mesh, which is then used for both training and validation.

#### 2.5.2. Training Data Augmentation

Only six dendritic branches are available for training, so we expanded the dataset by applying a series of geometric transformations. First, we computed the PCA of each mesh and projected its vertices onto the corresponding PCA plane. We then applied random scaling to the vertices in two separate steps, introduced random rotations in two additional steps, and added random translations. These transformations produced four augmented versions of each original mesh, resulting in a total of 30 dendritic branches available for training and validation. Of these, 22 branches were used for training, and the remaining 8 were used for validation.

#### 2.5.3. Testing dataset

For testing, we used data from the axon–spine coupling study, which includes a complete nanoconnectomic 3D reconstruction of hippocampal neuropil obtained via serial EM [20]. This dataset contains four independent annotations–two performed by each of two annotators [3]–providing detailed segmentation of dendritic spines. Out of 151 meshes, we selected 28 spiny dendritic branches for testing based on annotation consensus. Specifically, we included only those meshes where two or more annotators agreed on the presence of at least one dendritic spine. Additionally, for each spine mesh identified by the annotators, we retained only those meshes that were consistently labeled as spines by at least two of the selected annotations. This ensured that the testing set reflected a high-confidence subset of spiny dendritic structures.

It is important to note that some structure exhibiting spine-like morphology were not annotated as spines in the test set for two reasons. First, biologically, the presence of a synaptic area in the spine head is essential for defining a spine. Thus, even if a mesh appears spiny, it is not considered a spine if no synapse is present. Second, some meshes were excluded due to incomplete reconstruction–parts of the spine may have been cut off or lost during imaging.

In contrast, the training dataset does not apply as strict criteria for spine identification as the test set. As a result, the testing dataset introduces additional segmentation challenges that are not reflected in the training data. Our algorithm does not explicitly account for these differences, which may influence performance.

#### 2.5.4. Independent dataset

In addition to the training and testing datasets, we also used dendritic segments from the large-scale nanoconnectomic EM reconstruction of mouse neocortex by Kasthuri et al. [25]. This dataset provides densely reconstructed neuropil at nanometer resolution but does not include manual spine or shaft annotations. We therefore use it solely for qualitative visualization (Figure 7) to demonstrate that our method generalizes to independently acquired EM volumes. These data are not used for training, validation, or quantitative evaluation.

## 3. Results

This section presents a comprehensive evaluation of our algorithm’s performance in segmenting dendritic shafts and spines. We first describe the identification of dendritic shafts using geometric properties and a neural network-based approach. We then assess the ability of the algorithm to segment dendritic spines, highlighting both its strengths and limitations. Figure 7 illustrates the predicted dendritic shafts and spines from datasets from Kasthuri et al. [25], demonstrating that our method successfully identifies spines along the dendritic shaft by leveraging geometric properties such as Gaussian and mean curvature. However, misclassification can occur when the dendritic structure deviates significantly from the idealized cylindrical shape, underscoring the need for further improvements to the method.

To make these methods widely accessible, we have posted an open-source code repository on GitHub: [GitHub:curvature-based-dendrite-segmentation](https://github.com/aka-gera/curvature-based-machine-learning-for-automated-segmentation-of-dendritic-spines).

### 3.1. DNN Prediction Results Analysis

In this section, we present and analyze the prediction results of the three DNNs, **DNN**_1_, **DNN**_2_, and **DNN**_3_. The methodology is outlined in Section 2.2.4, and the evaluation here focuses on four key metrics: the training loss, the DICE coefficient (Dice Similarity Coefficient), the Jaccard index IoU (Intersection over Union), and the AUC (Area Under the ROC Curve, where ROC denotes the Receiver Operating Characteristic).

#### 3.1.1. Analysis of the performance

The training loss curves for the three models are shown in Figure S4. Among the three models, **DNN**_2_ achieves the most effective convergence, reaching a minimum loss of approximately 2.44, closely followed by **DNN**_3_ with a minimum loss of about 3.56. In contrast, the baseline architecture **DNN**_1_ shows the weakest convergence, with a substantially higher final loss of 6.73 throughout training. These results highlight the importance of the additional features and architectural refinements incorporated into **DNN**_2_ and **DNN**_3_ for improving optimization stability.

To quantitatively assess segmentation accuracy, we computed the IoU, DICE, and AUC scores between predicted and annotated vertices for the dendritic shaft and spines on both the training and test sets. The performance curves are presented in Figure S5. For **DNN**_2_, the IoU converges to average values of approximately 0.70 for the shaft and 0.81 for the spines, with corresponding DICE scores of about 0.88 and 0.92, respectively. **DNN**_3_ achieves comparable performance across both IoU and DICE metrics, whereas **DNN**_1_ lags behind with average IoU values of 0.55 (shaft) and 0.75 (spines), along with lower DICE scores overall. The AUC curves further support these trends: both **DNN**_2_ and **DNN**_3_ maintain consistently high AUC values throughout training, while **DNN**_1_ again shows weaker discriminative performance. Importantly, the similarity between training and testing curves across all metrics indicates good generalization and minimal overfitting.

Overall, these results demonstrate a clear progression in performance from **DNN**_1_ to **DNN**_3_. The incorporation of additional geometric and topological features in **DNN**_2_ and **DNN**_3_ substantially improves convergence and segmentation accuracy, underscoring the value of feature enrichment in dendritic spine detection.

### 3.2. Dendritic Spine Detection Analysis

Following dendritic shaft segmentation, we applied the spine detection algorithm described in Section 2.4 to identify spines. For each detected spine, we computed the IoU to evaluate segmentation accuracy. In addition, we calculated the IoU for the union of all detected spines to assess overall performance. We then computed accuracy, precision, recall, and F1-score to further quantify classification performance.

The segmentation results produced by the **DNN** models are shown in Figure 5. A clear qualitative progression is evident from **DNN**_1_ to the more advanced architectures **DNN**_2_ and **DNN**_3_. The **DNN**_1_ model frequently misclassifies portions of the shaft as spines, resulting in fragmented and noisy predictions. Incorporating additional geometric features—such as the distance to the shaft skeleton—allows **DNN**_2_ to reduce these errors and produce a more coherent separation between shaft and spine regions. Building on this, **DNN**_3_ integrates richer geometric and topological descriptors, enabling it to capture subtle curvature variations and resolve complex spine clusters more effectively. These improvements are visible not only in the 3D mesh renderings but also in the 2D slice, where the highlighted region clearly shows the reduction in mislabeling from **DNN**_1_ to **DNN**_3_. Overall, the predicted segmentations from **DNN**_3_ align most closely with the expert annotation. These qualitative trends are consistent with the quantitative results in Table S4, where both **DNN**_2_ and **DNN**_3_ outperform **DNN**_1_ across all evaluation metrics.

**Figure 5.**
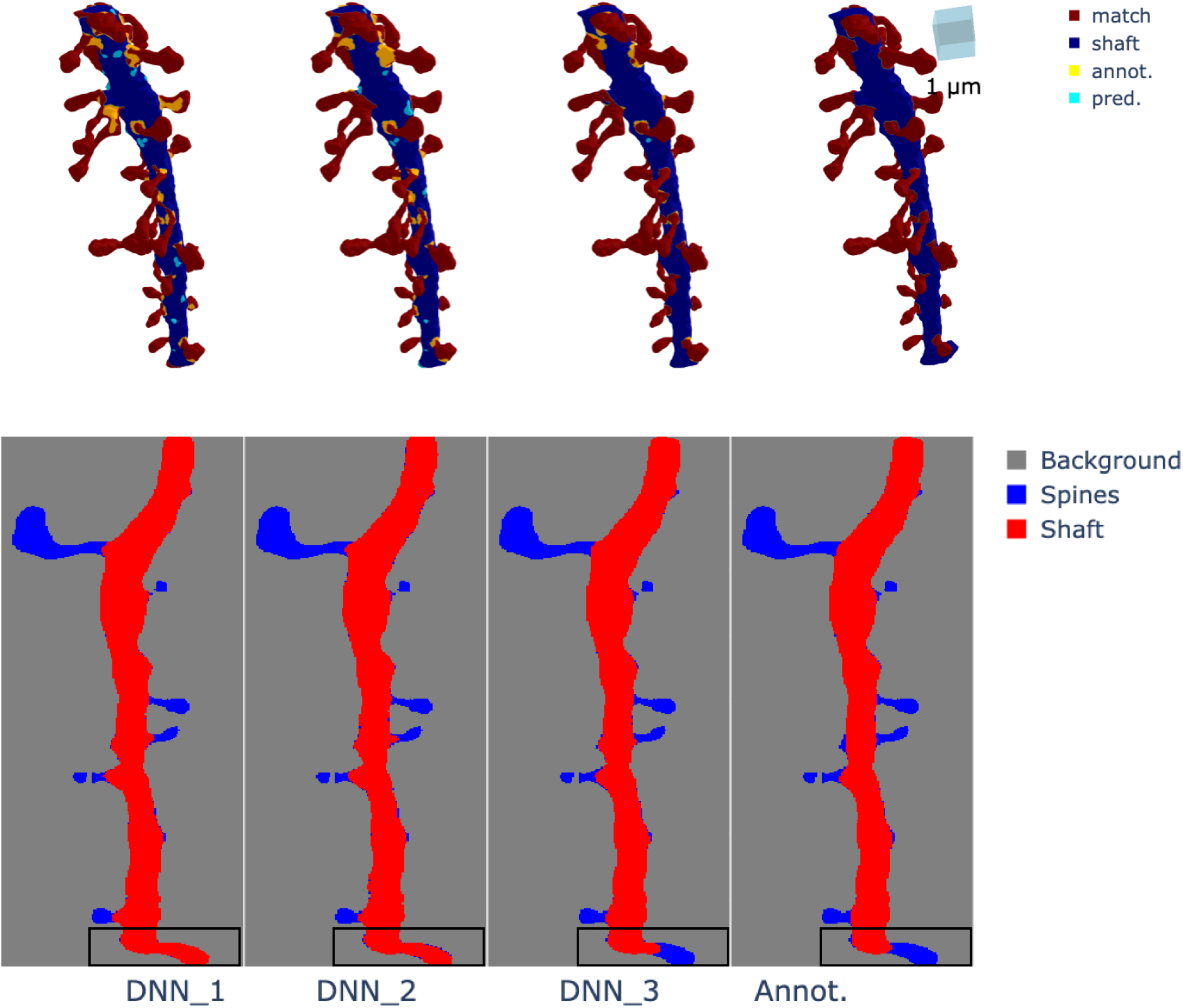
Comparison of segmentation results obtained using **DNN**_1_, **DNN**_2_, and **DNN**_3_. The first three columns correspond to the predictions from **DNN**_1_, **DNN**_2_, and **DNN**_3_, respectively, and the final column shows the expert spine annotation used as ground truth. The top row displays the 3D mesh representations, while the bottom row shows 2D slices of the same regions. In the meshes, red vertices indicate correct spine predictions that match the annotation, sky-blue vertices represent shaft regions misclassified as spines, and yellow vertices represent spine regions misclassified as shaft. **DNN**_1_ produces noisier results with more misclassifications, particularly in clustered regions. **DNN**_2_ reduces shaft-to-spine errors, and **DNN**_3_ achieves the most accurate segmentation overall, with fewer misclassifications and the closest agreement with the expert annotation. The slice highlights the region where mislabeling is most apparent, especially for **DNN**_1_ and **DNN**_2_. The triangular mesh is obtained from CA1 dendritic reconstructions [7–9].

#### 3.2.1. Quantitative Evaluation and Analysis

We evaluated segmentation performance using multiple complementary metrics, including AUC, Precision, Recall, and F1-score, computed under both IoU and DICE criteria, each with Standard and Union variants. These results, summarized in Table S4, enable a detailed comparison between our geometric-informed models (**DNN**_1_, **DNN**_2_, **DNN**_3_) and established CNN baselines (U-Net, VoxNet, VGG16FCN). Details on the training procedures and evaluation of the three CNN models are provided in Section S9. The weighting parameters used in the geometric-informed models are specified in (6), where the values of *a* for each model are listed and the parameter *b* is fixed at 0.81.

Among the CNN baselines, U-Net achieves the highest AUC (0.792), indicating strong global discriminative ability. However, AUC alone does not fully capture fine-scale segmentation quality. When examining Precision, Recall, and F1-score under the IoU and DICE criteria, the geometric-informed model **DNN**_3_ consistently delivers the strongest performance. Under the IoU–Union metric, **DNN**_3_ attains the highest Recall (0.873) and F1-score (0.845), indicating that it captures spine regions more completely than both the CNN baselines and the other geometric-informed variants. Its DICE–Union performance is even stronger, with an F1-score of 0.857 and a Recall of 0.895.

In contrast, **DNN**_1_ and **DNN**_2_ exhibit more limited recall, particularly under the Standard variants of IoU and DICE, where both models tend to miss a substantial fraction of true spines despite maintaining moderate-to-high precision. VoxNet and VGG16-FCN perform noticeably worse across all metrics, with lower AUC values (0.682 and 0.466, respectively) and substantially reduced F1-scores, especially under the IoU criterion. These results highlight the limitations of intensity-based CNN architectures when applied to thin, high-curvature structures such as dendritic spines.

The Union variants of IoU and DICE consistently yield higher Recall and F1-scores across all models, reflecting their ability to more fairly evaluate clustered or contiguous spine regions. This is particularly relevant in dense dendritic environments, where individual spine boundaries may be ambiguous. Under these more biologically meaningful metrics, **DNN**_3_ shows the clearest advantage, outperforming all baselines by a substantial margin. Overall, the quantitative results demonstrate a clear progression in performance from **DNN**_1_ to **DNN**_3_, with **DNN**_3_ providing the most reliable and complete segmentation of dendritic spines. The strong performance of **DNN**_3_ under both IoU and DICE—especially in the Union variants—underscores the value of incorporating geometric curvature cues for accurate spine–shaft segmentation.

### 3.3. Dendrite Spine Segmentation without Smoothing

In the segmentation of dendrite triangular meshes, curvature smoothing is typically required. However, when the number of vertices is large–more than 200,000–the process becomes computationally expensive, taking more than 20 hours to produce a smooth mesh. We note that by using **DNN**_3_, the computational time can be reduced by performing segmentation without smoothing, while still achieving a level of accuracy comparable to that obtained with smoothing.

We evaluated the effectiveness of curvature smoothing with **DNN**_3_ on both the training and test datasets and compared the accuracy. The computed accuracy for the test dataset is 0.658 (with and without smoothing), while for the training dataset it is 0.704 with smoothing and 0.780 without smoothing. These results show that smoothing the curvature of the triangular mesh when using **DNN**_3_ is unnecessary, and segmentation may even perform better without it.

This phenomenon may be due to the fact that the nine region-based segmentation features **S**^*k*^ enable the **DNN** to detect the neck area without requiring an enhanced curvature profile produced by smoothing. When we tested the same process on other **DNN** models, the accuracy was zero in both cases.

### 3.4. Application to Dendritic Surface Meshes with Many Vertices

We next use our algorithm to segment the spines of a dendritic mesh dataset containing a large number of vertices and multiple branches.

In particular, the dataset we study is from the reconstruction of a sub-volume of mouse neocortex [25]. We analyzed a dendritic mesh with 5,387,879 vertices and 10,777,005 faces. This dendritic mesh is about 13.42 times larger than the training dataset used in the previous sections (401,371 vertices and 802,750 faces). While most of the training and testing datasets contain only one main dendritic branch with attached spines, this dataset has nine branches with multiple spines. This highlights how much more complex this dataset is compared to the training and testing data.

Because this dataset is relatively large, the segmentation process differs slightly from that used in training. Computations were performed on a MacBook Pro with an Apple M4 Max chip and 64 GB of memory. When we attempted to segment the large dendritic mesh directly, the process failed due to computational constraints. Therefore, we reduced the size of the mesh using the simplification algorithm described in Section S6.2. The mesh was simplified to nearly the size of the largest training dataset, given (about 449,109 vertices and 900,000 faces), and we then proceeded with the same testing process as before.

The segmentation results are shown in Figure 7. Overall, **DNN**_3_ achieves the strongest performance, correctly labeling the majority of spines across the dendritic mesh. However, it still misclassifies a small region of the shaft as spines, as illustrated in the bottom panel (C). This error likely arises from local geometric similarities between the shaft surface and spine neck regions, particularly where the mesh bends or narrows.

In contrast, **DNN**_1_ also performs reasonably well and produces few large false-positive spine regions. However, closer inspection of panel (A) shows that **DNN**_1_ frequently misclassifies spine necks as shaft, and in some cases entire spines are labeled as shaft. These errors are consistent with the limited feature set used by **DNN**_1_, where Gaussian and mean curvature alone do not provide sufficient spatial context for reliable spine–shaft discrimination.

Finally, **DNN**_2_ exhibits the weakest performance on this dataset. It correctly identifies spines in only a subset of regions, while misclassifying large contiguous portions of the spines as shaft. This behavior suggests that the combination of shaft–skeleton distance, Gaussian curvature, and mean curvature features is insufficient to capture the complex local geometry present in this large-scale reconstruction.

### 3.5. Dendritic Spine Morphologic Parameters

In this section, we use the segmentation results obtained from our algorithm to compute several morphological parameters of the dendritic meshes in both the training and testing datasets. These parameters include the neck diameter, head diameter, and spine length.

The computation of head and neck diameters was performed using the spine skeleton. This skeleton was obtained by identifying the closest point of the triangular mesh skeleton to each vertex of the segmented spine mesh. To determine the neck and head diameters, we considered each segmented spine’s triangular mesh. For each vertex, we computed the shortest distance to its nearest skeleton point, as described in equation (5), while computing **D**. This procedure yielded a thickness profile along the spine. The neck radius was then defined as the minimum of these computed averages, starting from the skeleton vertices closest to the dendritic shaft. Conversely, the head diameter was defined as the maximum computed distance, measured from the skeleton vertices at the farthest point from the shaft. Additionally, to compute the length of the spine, we used the spine skeletonization and interpolated additional points along the skeleton using a spline technique. This interpolation increases the accuracy of the computed distances by smoothing and interpolating along the dendritic spine skeleton [13, 14, 53]. Additional details are provided in Section S7. The spine length was then computed as the sum of Euclidean distances between consecutive interpolated points:

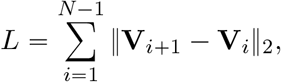

where **V**_*i*_ denotes the *i*-th interpolated vertex.

The distribution of morphological parameters in the training and large dendritic mesh datasets is shown in Figure 6. In the first column, the head diameter is plotted against the neck diameter of dendritic spines, while in the second column the spine volume is plotted against the spine area, with a colormap encoding spine length. For clearer visualization, some outliers were removed. Marginal histograms of head and neck diameters are also included to facilitate interpretation.

**Figure 6.**
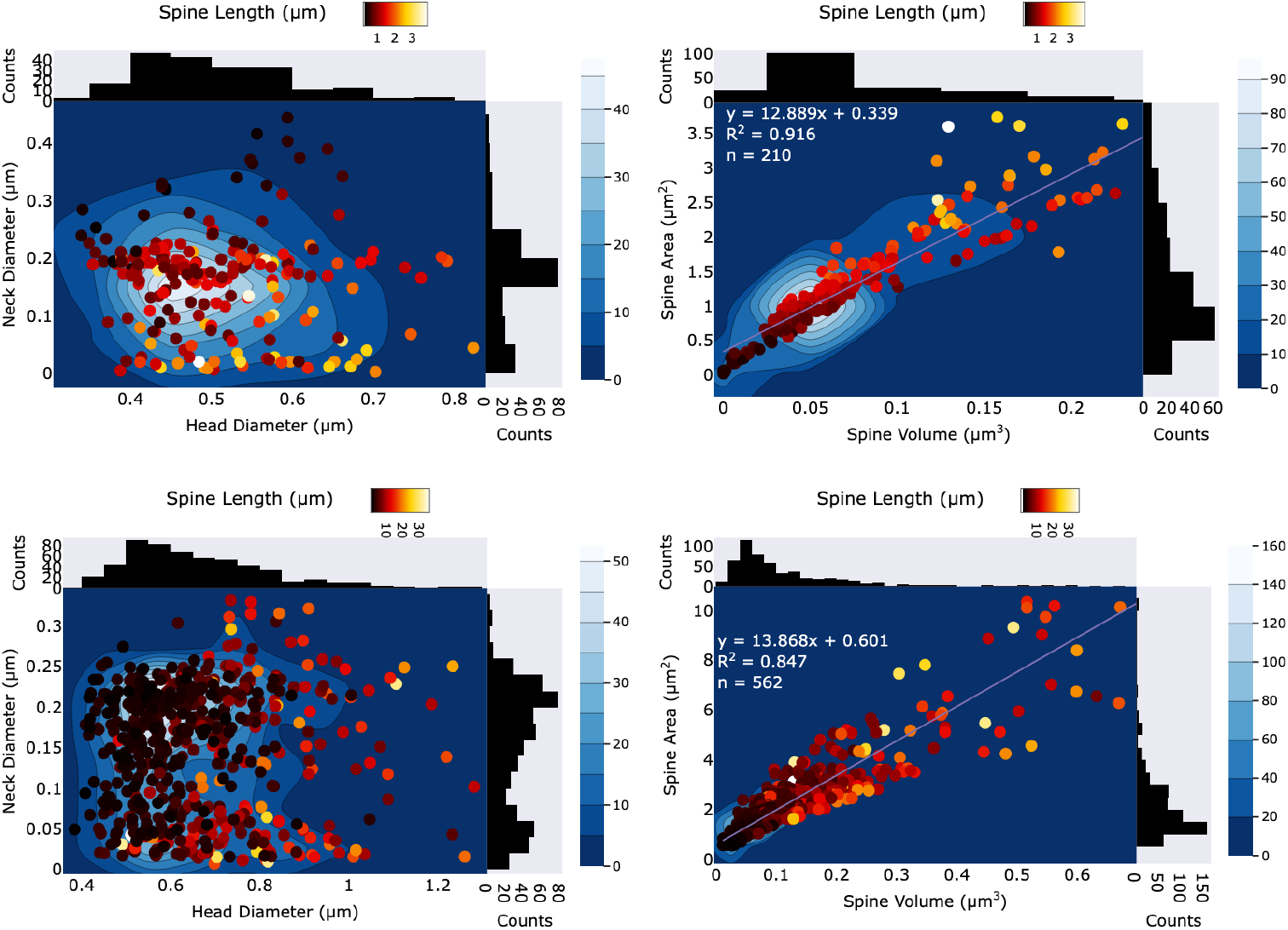
Distribution of morphological parameters in the training dataset and in the large dendritic mesh dataset, obtained using **DNN**_3_. In the first column, head diameter is plotted against neck diameter of dendritic spines, while in the second column spine volume is plotted against spine area. A colormap encodes spine length. The segmented training dataset contains 210 spines, with an average neck diameter of 0.166 ± 0.0969 *µ*m, an average head diameter of 0.512 ± 0.101 *µ*m, and an average spine length of 1.35 ± 0.694 *µ*m. The average spine volume is 0.122 ± 0.121 *µ*m^3^ and the average spine area is 2.30 ± 1.83 *µ*m^2^, yielding an average area-to-volume ratio of 12.9 with a coefficient of determination *R*^2^ = 0.916. The segmented large dendritic mesh dataset contains 562 spines, with an average neck diameter of 0.148 ± 0.0787 *µ*m, an average head diameter of 0.661 ± 0.164 *µ*m, and an average spine length of 5.57 ± 5.34 *µ*m. The average spine volume is 0.090 ± 0.085 *µ*m^3^ and the average spine area is 1.53 ± 1.15 *µ*m^2^, yielding an average area-to-volume ratio of 13.9 with a coefficient of determination *R*^2^ = 0.847. The dataset is derived from the Nanoconnectomic 3D EM reconstruction of hippocampal neuropil from the axon–spine coupling study [3].

Quantitatively, the segmented training dataset contains 210 spines, with an average neck diameter of 0.166 ± 0.0969 *µ*m, an average head diameter of 0.512 ± 0.101 *µ*m, and an average spine length of 1.35 ± 0.694 *µ*m. The average spine volume is 0.122 ± 0.121 *µ*m^3^ and the average spine area is 2.30 ± 1.83 *µ*m^2^, yielding an average area-to-volume ratio of 12.9. Importantly, the relationship between spine volume and area is very strong, as reflected by a high coefficient of determination (*R*^2^ = 0.916).

In contrast, the segmented large dendritic mesh dataset contains 562 spines, with an average neck diameter of 0.148 ± 0.0787 *µ*m, an average head diameter of 0.661 ± 0.164 *µ*m, and an average spine length of 5.57 ± 5.34 *µ*m. The average spine volume is 0.090 ± 0.085 *µ*m^3^ and the average spine area is 1.53 ± 1.15 *µ*m^2^, yielding an average area-to-volume ratio of 13.9. Here too, the relationship between spine volume and area remains strong, with a coefficient of determination of *R*^2^ = 0.847.

## 4. Discussion

Despite its overall promise, our algorithm encounters challenges in segmenting spines with complex morphologies and spatial arrangements. Below, we discuss three key failure cases, illustrated in Figures 8. The consistently high precision suggests that when a spine is detected, it is almost always correct (few false positives). However, the moderate recall for IoU alone indicates that some spines are missed (false negatives), potentially affecting downstream morphological analyses. The substantial performance improvement with union-based IoU suggests that a more inclusive segmentation strategy enhances detection reliability.

**Figure 7.**
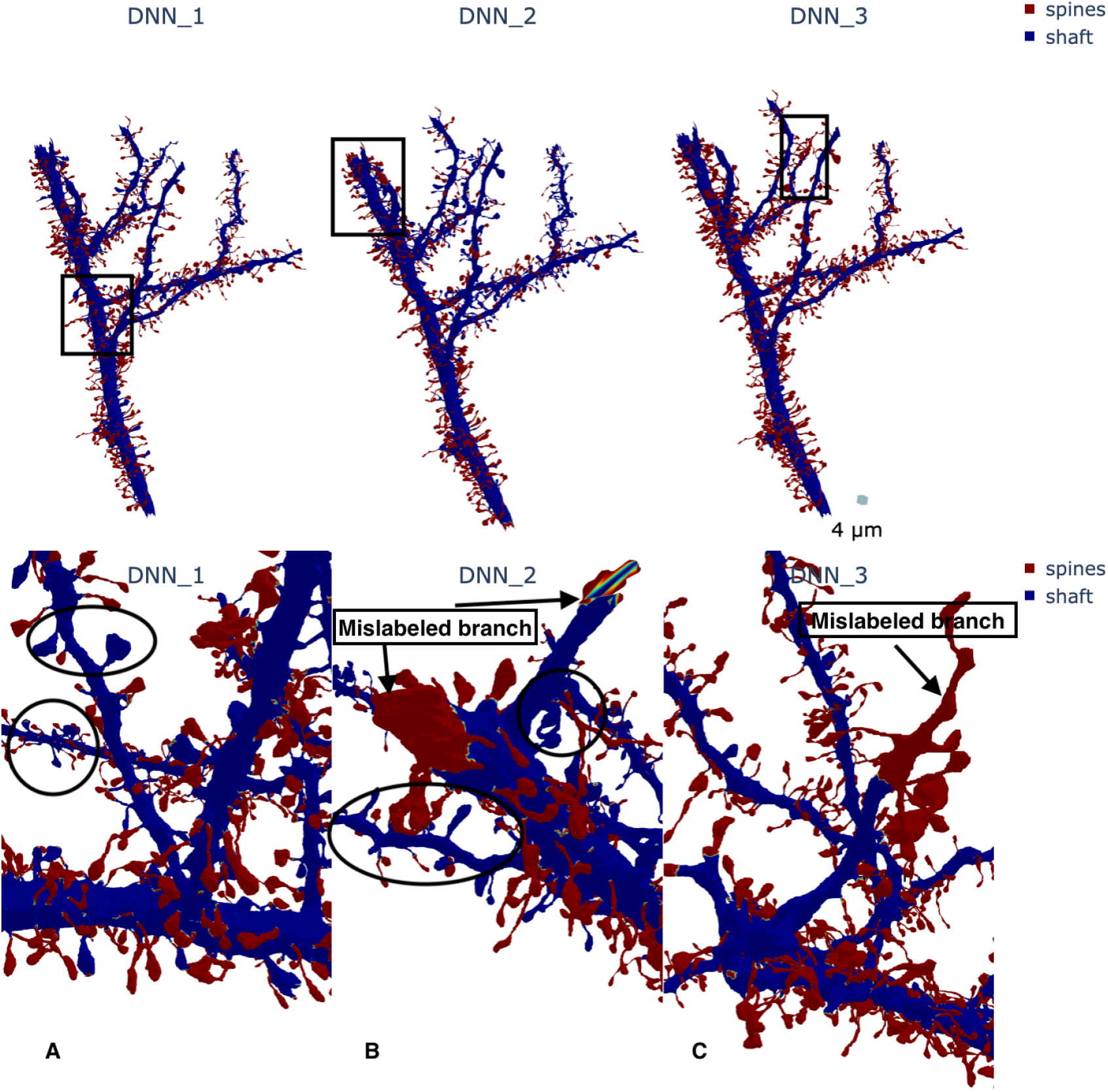
Comparison of segmentation results obtained using **DNN**_1_, **DNN**_2_, and **DNN**_3_ on a large-scale nanoconnectomic EM reconstruction of mouse neocortex from Kasthuri et al. [25]. In all panels, blue indicates the dendritic shaft mesh, while red highlights the segmented spines. The top panel shows the full dendritic segment with predictions from the three models. All models correctly identify the majority of spines, but notable differences emerge upon closer inspection. Three boxed regions highlight areas of disagreement, which are shown in detail in the bottom panels. Panel (A) shows spine meshes that **DNN**_1_ incorrectly labels as shaft. Panel (B) illustrates dendritic sub-branches that **DNN**_2_ misclassifies as spines. Panel (C) shows a similar misclassification pattern for **DNN**_3_, where local curvature and branching geometry lead to spurious spine predictions.

**Figure 8.**
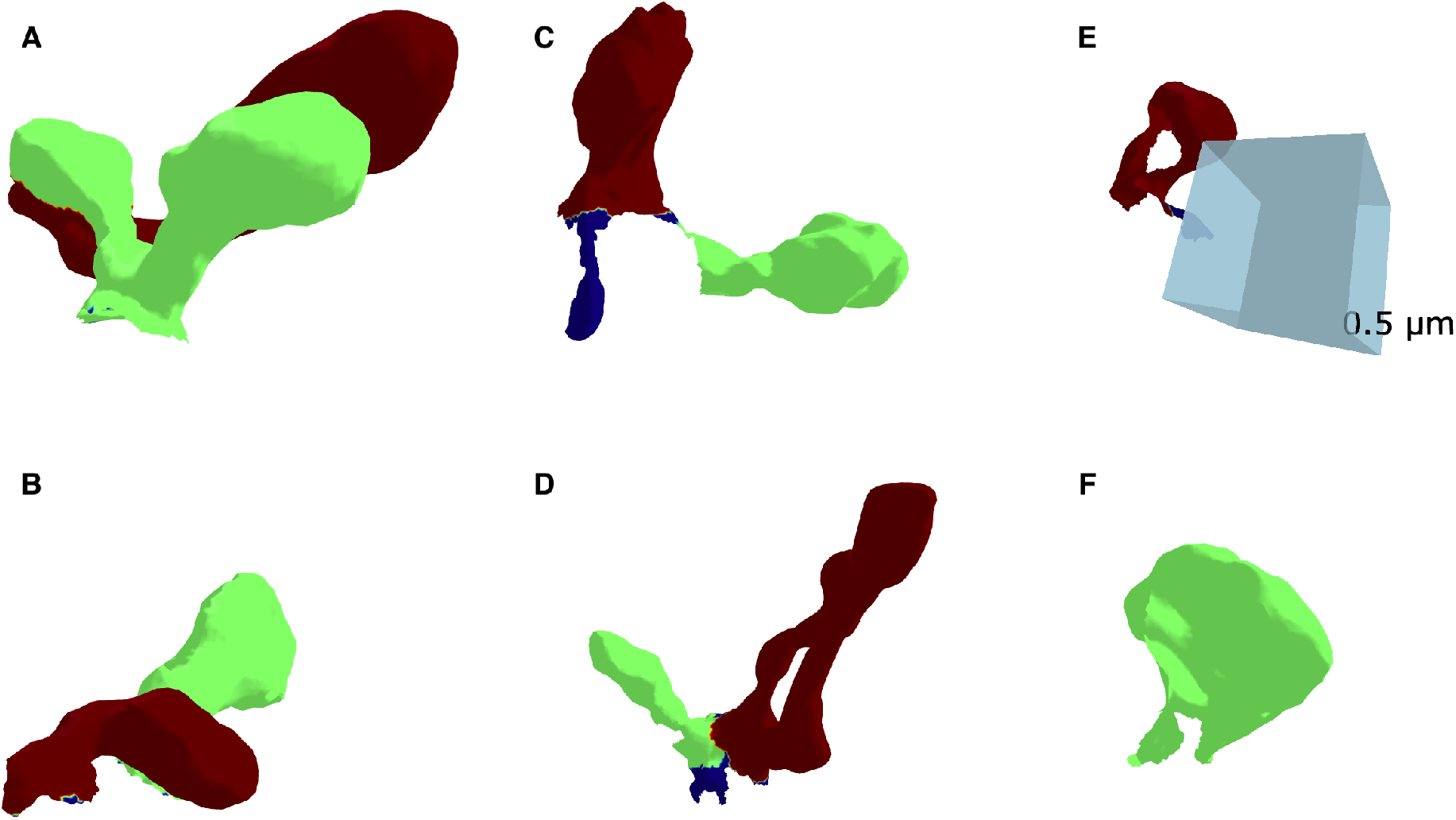
Examples of misclassification encountered during spine segmentation. In the figure, blue indicates the shaft mesh, while other colors indicate spine meshes that were misclassified. Panels (A–D) illustrate cases where multiple spines were incorrectly merged and classified as a single spine. Panel (E) depicts a region of the dendritic shaft that was mistakenly labeled as a spine, while Panel (F) shows a fraction of a spine that was only partially labeled. These examples highlight the challenges of accurately separating spines in dense clusters. The triangular mesh is obtained from CA1 dendritic reconstructions [7–9].

### 4.1. Limitations in Spine Group Segmentation

Dendritic spines frequently appear in dense clusters, complicating their segmentation. These spine groups are two or more spines whose vertices are directly connected, and after the shaft is removed they still group together. Figure 8 shows a region containing tightly packed spines where the algorithm struggles to distinguish individual structures due to overlapping curvature features. This limitation highlights the need for improved clustering-based segmentation techniques or the integration of additional geometric descriptors capable of differentiating individual spines in high-density regions.

#### 4.1.1. Misclassification of Dendritic Shaft Regions as Spines

Another failure mode involves the erroneous classification of dendritic shaft regions as spines, as depicted in Figure 8[F]. This misclassification likely stems from local curvature variations that resemble spine-like features. To mitigate this issue, we propose the incorporation of spatial continuity constraints, contextual neighborhood information, or multi-scale curvature analysis to better distinguish dendritic shafts from true spine structures.

#### 4.1.2. Scalability and Large Dataset Limitations

A further limitation arises from the scale of the datasets used in training and evaluation. High-resolution 3D EM reconstructions of dendritic segments generate extremely large meshes, often containing millions of vertices. Processing such datasets requires substantial computational resources, both in terms of memory and runtime. This constraint limits the feasibility of applying our method to very large-scale reconstructions or to entire brain regions without significant preprocessing or downsampling. Moreover, the need to balance mesh resolution with computational efficiency may lead to the loss of fine structural details, particularly in thin spine necks or small protrusions. Addressing this limitation will require the development of more efficient algorithms, parallelized implementations, or hierarchical multi-resolution approaches that can scale to increasingly large datasets while preserving biologically relevant detail.

## 5. Conclusion

In this work, we developed and evaluated three geometric-informed deep neural network architectures for dendritic shaft and spine segmentation. Our results demonstrate a clear progression in performance from the baseline **DNN**_1_ to the enhanced models **DNN**_2_ and **DNN**_3_, with **DNN**_3_ achieving the strongest overall segmentation accuracy. By incorporating curvature-based and topology-aware features, these models substantially improve both training stability and predictive performance, particularly in regions with dense spine clusters or complex branching patterns.

A key advantage of the geometric-informed approach is its computational efficiency relative to volumetric 3D CNNs. CNN-based models require dense voxel representations that scale cubically with resolution, making them memory intensive and slow when applied to high-resolution EM reconstructions. In contrast, our geometric models operate directly on mesh-derived features, avoiding voxelization and enabling faster inference on large or irregular geometries. As a result, **DNN**_3_ not only surpasses all CNN baselines in accuracy but also scales more effectively to high-resolution meshes, making it the preferred choice for detailed EM datasets where preserving fine structural detail is essential.

Overall, our findings highlight the potential of feature-enriched geometric learning for robust dendritic spine detection. By improving segmentation accuracy, computational efficiency, and scalability, these models provide a strong foundation for downstream morphological analyses and quantitative studies of synaptic connectivity. Continued refinement of geometric descriptors, multi-resolution strategies, and post-processing methods will further enhance the reliability and applicability of this framework to increasingly large and complex neural reconstructions.

## Acknowledgments

We acknowledge Myles Joyce for help with spine segmentation. This work was supported by NSF Technology Hub #1707356, NSF NeuroNex2 #2014862, NSF award 2219894, and NIH grants R01MH095980 and R56MH139176 to KMH and NIH grant T32 NS007292 to AKAG. We acknowledge use of the Brandeis High Performance Computing Cluster (HPCC) which is partially supported by the NSF through DMR-MRSEC 2011846 and OAC-1920147.

