## Supplemental Information for "Curvature-based machine learning method for automated segmentation of dendritic spines"

### Supplemental Materials and Methods

#### S1. Enhancement of Curvature through Image Processing

For the first panel of Figure 3, we choose parameters that emphasize the neck region of the triangular dendritic mesh. In particular, we visualize the function

$$F = \widetilde{\mathbf{H}} + \widetilde{\mathbf{K}} \qquad \begin{cases} a_{\mathbf{H}} & = 5 \\ b_{\mathbf{H}} & = 10 \\ a_{\mathbf{K}} & = -1 \\ b_{\mathbf{K}} & = -10 \end{cases} .$$

This combination enhances the contrast of the neck surface and provides an interpretable geometric baseline for comparison with the learned logits.

#### S2. Training Details

The **DNN** architecture was built using `TensorFlow` (v2.16.2) [1], and training was performed for more than 1500 epochs for each model, after which the loss had converged. We trained the network using the Adam optimizer [28] with a piecewise constant learning rate schedule. The learning rate was set to $10^{-2}$ for the first 1000 training steps, reduced to $10^{-3}$ for steps 1000–3000, and further reduced to $5 \times 10^{-4}$ for the remainder of training. Only a single batch was used per training step, corresponding to a batch size of 1, because each dendritic mesh is processed independently and cannot be meaningfully combined with others within the same batch.

L1 and L2 regularization were applied to the hidden layers to improve generalization and reduce overfitting. This dual regularization strategy encourages sparsity in the learned weights while penalizing large parameter values [21,49], thereby improving robustness across diverse inputs.

The class imbalance between shaft and spine vertices was handled using class weights derived from the inverse log-frequency of each class. Let $c_k$ denote the number of vertices belonging to class $k$ and $T = \sum_k c_k$ the total number of vertices. The unnormalized class weight is

$$w'_k = \max\left(\log\left(\frac{2T}{c_k}\right), 1\right),$$

and the final normalized weights were obtained by $w_k = w'_k / \sum_j w'_j$. These weights were applied to the cross-entropy loss during training to prevent the model from being biased toward the majority class.

#### S3. Computer Specifics

All geometric models ($\mathbf{DNN}_1$, $\mathbf{DNN}_2$, $\mathbf{DNN}_3$) were trained on an HPCC CPU-only node equipped with 2×Intel Cascade Lake CPUs (40 total cores) and 754 GB of RAM. The CNN-based models (U-Net, VoxNet, VGG16–FCN) could not be trained on a local machine due to their substantially higher computational and memory requirements. Even after downsampling the meshes to 5,000 vertices, the CNN architectures exceeded the memory capacity of the local system and were therefore trained exclusively on the HPCC node.

All testing and inference experiments were performed on a MacBook Pro equipped with an Apple M4 Max chip and 64 GB of unified memory.

The geometric models run an order of magnitude faster than the CNN baselines, with $\mathbf{DNN}_3$ achieving the lowest runtime (43–45 s/iteration), while the CNN models require 5–8 times longer per iteration despite comparable memory usage. Table S2 summarizes the runtime and memory characteristics of all evaluated models.

### S4. Performance Metrics

Various performance metrics were selected to evaluate the quality of the predicted segmentations. The DICE coefficient, Jaccard index (IoU), and AUC provide complementary perspectives on segmentation performance. The DICE coefficient quantifies the overlap between the predicted region $P$ and the ground-truth region $G$, and is defined as

$$\mathrm{DICE} = \frac{2|P \cap G|}{|P| + |G|}.$$

Higher values indicate stronger agreement between prediction and ground truth. The Jaccard index (IoU) offers a stricter measure of overlap, computed as

$$\mathrm{IoU} = \frac{|P \cap G|}{|P \cup G|},$$

and penalizes both false positives and false negatives. Finally, the AUC (Area Under the ROC Curve) evaluates the model's ability to discriminate between classes across all possible thresholds; the ROC (Receiver Operating Characteristic) curve plots the true-positive rate against the false-positive rate. Together, these metrics capture spatial accuracy, overlap quality, and classification discriminability, providing a comprehensive assessment of segmentation performance.

### S5. Discrete Gaussian and Mean Curvature

In this section, we briefly review the discrete differential geometry formulation presented in [36, 40, 48, 55] to derive the Gaussian and the mean curvature of triangular meshes. Assume the mesh has $\mathcal{V}$ vertices and $\mathcal{T}$ triangular faces. Each triangular face $\mathcal{T}_l \in \mathcal{T}$ has vertices denoted by $k_i^l$, $k_{i+1}^l$, and $k_{i+2}^l$. These vertices are indexed in a counterclockwise order around the face, defined as follows:
Let $k_{\text{next}}(\cdot)$ denote the next vertex in the counterclockwise direction from $\cdot$. Then:

- $k_i^l$ is the first vertex.
- $k_{i+1}^l = k_{\text{next}}(k_i^l)$ is the next vertex in the counterclockwise direction.
- $k_{i+2}^l = k_{\text{next}}(k_{i+1}^l)$ is the vertex following $k_{i+1}^l$ in the counterclockwise direction. Then $k_{i+2}^l = k_{\text{next}}(k_{\text{next}}(k_i^l))$ .
- $k_i^l = k_{\text{next}}(k_{i+2}^l) = k_{\text{next}}(k_{\text{next}}(k_{\text{next}}(k_i^l)))$ completes the cycle.

This leads to the vertex indices cycling according to:

$$k_i^l = k_{(i-1 \mod 3)+1}^l,$$

where $(i \mod 3) + 1$ cycles through the indices $\{1, 2, 3\}$. Moreover, we will denote the set of indices of the 1-ring neighbors of a vertex with index $k$ by $\mathcal{N}_k$. This set includes the indices of the vertices whose edges are connected to the vertex $\mathbf{X}_k$:

$$\mathcal{N}_k = \{j \mid \text{there exists an edge between } \mathbf{X}_k \text{ and } \mathbf{X}_j\}.$$

S5.1. **Discrete Gaussian Curvature.** Let us consider an infinitesimal area $\mathcal{A}$ and denote its diameter by diam($\mathcal{A}$). Additionally, let $\mathcal{A}^{\mathcal{G}}$ denote the area of the image of the Gauss map associated with $\mathcal{A}$. We can express the discrete Gaussian curvature, $\widehat{\kappa}_G$ at a vertex $p = \mathbf{X}_l$ as:

$$\widehat{\kappa}_G = \frac{1}{\mathcal{A}} \int\int_{\mathcal{A}} \kappa_G dA = \frac{1}{\mathcal{A}} \sum_{p \in \mathcal{A}} \mathbf{K}_p, \text{ with } \mathbf{K}_p = 2\pi - \sum_{j \in \mathcal{N}_i} \theta_j \tag{7}$$

where $\theta_j$ are the interior angles at $p = \mathbf{X}_i$ of the triangles meeting there. The term $\mathbf{K}_p$ is known as the defect angle at $p$. Considering vertices $\mathbf{X}_j$ in the 1-ring neighborhood of $p = \mathbf{X}_l$, see Fig S1, the angle $\theta_j$ can be computed as:

$$\cos\theta_j = \frac{(\mathbf{X}_{j-1} - \mathbf{X}_l) \cdot (\mathbf{X}_j - \mathbf{X}_l)}{\|(\mathbf{X}_{j-1} - \mathbf{X}_l) \cdot (\mathbf{X}_j - \mathbf{X}_l)\|} = \frac{\mathbf{E}_{l,j-1} \cdot \mathbf{E}_{ij}}{\|\mathbf{E}_{l,j-1} \cdot \mathbf{E}_{ij}\|},$$

where $\mathbf{E}_{l,j} = \mathbf{X}_j - \mathbf{X}_l$ is the edge of the vertices $\mathbf{X}_i, \mathbf{X}_j$.

S5.2. **Mean curvature.** The discrete mean curvature, $\widehat{\kappa}_H$, at a vertex $p$ is defined as:

$$\widehat{\kappa}_H = \frac{1}{\mathcal{A}} \int\int_{\mathcal{A}} \kappa_H dA = \frac{1}{\mathcal{A}} \sum_{p \in \mathcal{A}} \mathbf{H}_p, \qquad 2\mathbf{H}_p = \int\int_{\mathcal{A}} \kappa_H dA. \tag{8}$$

To compute $\mathbf{H}_p$, we consider two equivalent methods [36,40,48,55]. First, assuming vertices adjacent to $p$ in cyclic order are $\mathbf{X}_j, \mathbf{X}_{j+1}, \ldots \mathbf{X}_{j+n}$. we have:

$$\begin{aligned} 2\mathbf{H}_p &= \sum_{m=j} \mathbf{E}_{l,m} \times \mathbf{n}_{m+1} - \mathbf{E}_{l,m} \times \mathbf{n}_m \\ &= - \sum_{m=j} \mathbf{E}_{l,m} \times \mathbf{n}_m, \end{aligned}$$

where $\mathbf{n}_j$ is the normal vector to the triangle with vertices $\mathbf{X}_l, \mathbf{X}_{j-1}, \mathbf{X}_j$:

$$\mathbf{n}_j = \frac{(\mathbf{X}_{j-1} - \mathbf{X}_l) \times (\mathbf{X}_j - \mathbf{X}_l)}{\|(\mathbf{X}_{j-1} - \mathbf{X}_l) \times (\mathbf{X}_j - \mathbf{X}_l)\|} = \frac{\mathbf{E}_{l,j-1} \times \mathbf{E}_{l,j}}{\|\mathbf{E}_{l,j-1} \times \mathbf{E}_{l,j}\|}.$$

Alternatively, the second method involves the integral of the Laplace-Beltrami operator [36]:

$$2\mathbf{H}_p = \sum_{j \in \mathcal{N}_i} (\cot\alpha_{l,j} + \cot\beta_{l,j})(\mathbf{X}_l - \mathbf{X}_j),$$

where $\alpha_{l,j}$ and $\beta_{l,j}$ are the angles opposite edge $\mathbf{E}_{l,j}$ in the two incident triangles, as shown in Fig S1.

S5.3. **Vector Area.** In this section, we compute the area $\mathcal{A}$ as in (7), (8), the sum of all the regions that contain $p$,

$$\mathcal{A} = \sum_{p \in \mathcal{A}} A_p.$$

To do this, $A_p$ is computed using either the conic area or the barycentric formula that we will describe below. Both methods are equivalent in any case [36].

S5.3.1. *Conic area.* The formula used to compute the vector area is:

$$\mathbf{A} = \frac{1}{2}\int\int_{\mathcal{A}} \mathbf{n} dA,$$

and specifically in the triangle with edge $\mathbf{E}_i, \mathbf{E}_j$ case, the area is given by:

$$A_{ij} = \frac{1}{2}\left\|\mathbf{E}_i \times \mathbf{E}_j\right\|$$

As the conic area is equal to the third of the area in the 1-ring neighborhood of $p$, we have:

$$A_p = \frac{1}{3}A = \frac{1}{6}\sum_{j\in\mathcal{N}_1(i)} \left\|\mathbf{E}_{i,j-1} \times \mathbf{E}_{ij}\right\|$$

S5.3.2. *Barycenter Area.* To compute the barycentric area (represented in blue in Figure S1), we first find the centroid $\mathbf{C}_{ij}$ of each triangle in the 1-ring neighborhood:

$$\mathbf{C}_{ij} = \frac{1}{3}(\mathbf{X}_i + \mathbf{X}_{j-1} + \mathbf{X}_j).$$

Next, we compute the middle point $\mathbf{X}_{ij}$ of each edge $\mathbf{X}_i, \mathbf{X}_j$:

$$\mathbf{M}_{ij} = \frac{1}{2}(\mathbf{X}_i + \mathbf{X}_j).$$

Then the barycentric area corresponding to the triangle with the vertices $\mathbf{X}_i, \mathbf{X}_{j-1}, \mathbf{X}_j$ is

$$A_{ij} = \frac{1}{2}\left\|(\mathbf{M}_{i,j-1} - \mathbf{C}_{ij}) \times (\mathbf{X}_i - \mathbf{C}_{ij})\right\| + \frac{1}{2}\left\|(\mathbf{M}_{ij} - \mathbf{C}_{ij}) \times (\mathbf{X}_i - \mathbf{C}_{ij})\right\|.$$

Thus, the barycentric area of the 1-ring neighborhood $A_p$ is:

$$A_p = \sum_{j\in\mathcal{N}_1(i)} A_{ij}.$$

### S6. Skeletonization

In this section, we describe the skeletonization algorithm applied to a 3D mesh using Python packages such as `trimesh`, `open3d`, and `scikit-image`. All experiments were performed in a Python 3.9.6 environment with the following package versions: `numpy` (v2.2.6) [18], `open3d` (v0.18.0) [61], `trimesh` (v4.6.8) [12], and `scikit-image` (v0.24.0) [51]. The goal of this algorithm is to extract a simplified, one-voxel-wide medial axis from a complex dendritic mesh, which can then be used for further geometric analysis and as input to the DNN.
To perform skeletonization, the first step is to ensure that the mesh is watertight. A watertight mesh is a closed surface with no gaps, holes, or disconnected edges, which guarantees a well-defined interior and exterior. If the input mesh is not watertight, we apply a wrapping procedure based on Poisson surface reconstruction to generate a closed representation. Once the watertight mesh is obtained, the pipeline proceeds through mesh simplification, voxelization, skeleton extraction, and vertex-to-skeleton mapping.

S6.1. **Mesh Wrapping.** To ensure the mesh is watertight and suitable for skeletonization, we apply a wrapping procedure based on Poisson surface reconstruction [26, 27, 61]. This step converts the input mesh into a uniformly sampled point cloud, estimates and orients normals, and then reconstructs a closed surface. The implementation is shown below:

```
mesh = trimesh.Trimesh(vertices=vertices, faces=faces)
o3d_mesh = o3d.geometry.TriangleMesh()
o3d_mesh.vertices = o3d.utility.Vector3dVector(mesh.vertices)
o3d_mesh.triangles = o3d.utility.Vector3iVector(mesh.faces)

# Sample points uniformly from the surface
pcd = o3d_mesh.sample_points_poisson_disk(
    number_of_points=number_of_points
)

# Estimate and orient normals
pcd.estimate_normals(
    search_param=o3d.geometry.KDTreeSearchParamHybrid(
        radius=radius, max_nn=max_nn
    )
)
pcd.orient_normals_consistent_tangent_plane(k=10)

# Reconstruct a watertight mesh using Poisson surface reconstruction
mesh_poisson, _ = o3d.geometry.TriangleMesh.create_from_point_cloud_poisson(
    pcd, depth=8
)

vertices = np.asarray(mesh_poisson.vertices)
faces = np.asarray(mesh_poisson.triangles)
```

Here, we first convert the input vertices and faces into a `trimesh.Trimesh` object and then into an Open3D `TriangleMesh`. From this mesh, we generate a uniformly sampled point cloud using Poisson disk sampling [58, 61] Normals are estimated and consistently oriented to ensure correct surface reconstruction. Finally, Poisson surface reconstruction produces a watertight mesh, which is returned as arrays of vertices and faces for subsequent processing. To provide an alternative geometry-preserving option, we also include an implementation of the `alpha-wrap` method, which is based on the classical $\alpha$-shape formulation [15] and implemented in `PyMeshLab` [11]. This approach generates a watertight surface by constructing an $\alpha$-shape envelope around the input mesh. The corresponding code is shown below:

```
ms = pymeshlab.MeshSet()
ms.add_mesh(
    pymeshlab.Mesh(
        vertex_matrix=vertices,
        face_matrix=faces
    )
)
```

```
# Apply alpha-wrap reconstruction
ms.apply_filter(
    'generate_alpha_wrap',
    alpha=pymeshlab.PercentageValue(alpha_fraction),
    offset=pymeshlab.PercentageValue(offset_fraction)
)

mesh_wrap = ms.current_mesh()
vertices_wrap = mesh_wrap.vertex_matrix()
faces_wrap = mesh_wrap.face_matrix()
```

Here, the input vertices and faces are loaded into a `MeshSet`, and the `generate_alpha_wrap` filter constructs a watertight surface based on the specified `alpha` and `offset` parameters. The resulting mesh is then extracted as arrays of wrapped vertices and faces for downstream processing.

S6.2. **Mesh Simplification.** We apply a mesh simplification algorithm to reduce the size of the mesh in cases where the number of vertices is very large. This step is crucial for lowering computational complexity and removing unnecessary geometric detail that may interfere with accurate skeletonization. The simplification is performed using quadric decimation [17, 61] in Open3D, as shown in the following code:

```
mesh = trimesh.Trimesh(vertices=vertices, faces=faces)
o3d_mesh = o3d.geometry.TriangleMesh()
o3d_mesh.vertices = o3d.utility.Vector3dVector(mesh.vertices)
o3d_mesh.triangles = o3d.utility.Vector3iVector(mesh.faces)

# Simplify the mesh using quadric decimation
o3d_mesh = o3d_mesh.simplify_quadric_decimation(
    target_number_of_triangles=target_number_of_triangles
)

# Convert back to a trimesh object for compatibility
mesh = trimesh.Trimesh(
    vertices=np.asarray(o3d_mesh.vertices),
    faces=np.asarray(o3d_mesh.triangles)
)
```

Here, we first create a `trimesh.Trimesh` object from the input vertices and faces. This mesh is then converted into an Open3D `TriangleMesh`, which supports advanced mesh processing operations. The `simplify_quadric_decimation` method reduces the number of triangles while preserving the overall geometry and removing fine details. Finally, the simplified mesh is converted back into a `trimesh` object for compatibility with subsequent steps in the pipeline.

S6.3. **Voxelization.** Next, we voxelize the simplified mesh to convert it into a discrete 3D grid representation. This step enables morphological operations such as thinning and skeletonization. The voxelization process is controlled by a resolution parameter, which determines the granularity of the voxel grid:

```
pitch =(np.median(mesh.edges_unique_length) * 0.25
```

```
voxelized = mesh.voxelized(pitch=pitch))
filled = voxelized.fill(method="orthogonal")
voxels = filled.matrix.astype(bool)
```

The parameter `pitch` in `mesh.voxelized` is determined from the median edge length of the mesh, scaled by a factor of 0.25. This adaptive choice ties the voxel size to the geometric detail of the mesh, ensuring that the discretization captures fine structures without producing an excessively large grid. The `fill()` method is then applied to close internal cavities, yielding a watertight solid volume. Finally, the voxel grid is converted into a boolean array, where each entry indicates whether a voxel is occupied, providing a suitable representation for subsequent skeletonization.

### S7. Spline-Based Interpolation

Given the set of spine vertices, we apply the `splprep` function from the `scipy.interpolate` module in `scipy`(v0.24.0) [53], which computes a B-spline representation of an $N$-dimensional parametric curve. This enables smoothing and interpolation along the dendritic spine skeleton [13, 14]. The spline is then evaluated using `splev` to obtain a dense set of interpolated points along the curve:

```
points = spine_vertices.T
tck, u = splprep([points[0], points[1], points[2]],
                 s=spline_smooth,
                 k=max(1, min(3, len(points[0]) - 1)))
x_fine, y_fine, z_fine = splev(np.linspace(0, 1, line_num_points), tck)
interpolated_vertices = np.column_stack([x_fine, y_fine, z_fine])
```

Here, `line_num_points` controls the density of interpolation; in our experiments we set it to 150, and the smoothing factor `spline_smooth` was chosen as 0.03. The resulting interpolated vertices are ordered sequentially along the spline, which is ensured by the spline parameterization itself.

### S8. Confusion Matrix Analysis

We assess the performance of our model by computing standard classification metrics, including accuracy, precision, recall, and the F1-score. This section details the computation process.

To begin, we illustrate the confusion matrix, which provides a detailed comparison between the predicted classifications and the ground truth annotations. This allows for an in-depth evaluation of classification performance. In our framework, a group of vertices is predicted as spine if its IoU with the annotated spine vertices it overlaps exceeds 0.7.

The classification outcomes are defined as follows:

- **True Positives (TP)**: Spines correctly identified by the algorithm.
- **False Negatives (FN)**: Spines present in the ground truth but missed by the algorithm.
- **False Positives (FP)**: Shaft regions incorrectly classified as spines.
- **True Negatives (TN)**: Shaft regions correctly classified.

Once the confusion matrix is computed, we derive the following performance metrics:

Precision. Precision evaluates the reliability of spine predictions:

$$\text{Precision} = \frac{\text{TP}}{\text{TP} + \text{FP}}.$$

Recall. Recall measures the model's ability to detect actual spines:

$$\text{Recall} = \frac{\text{TP}}{\text{TP} + \text{FN}}.$$

F1-score. The F1-score provides a balanced measure of precision and recall:

$$\text{F1-score} = 2 \times \frac{\text{Precision} \times \text{Recall}}{\text{Precision} + \text{Recall}}.$$

### S9. Dendritic Spine Analysis Using 3D CNNs

In this section, we evaluate dendritic spine segmentation using several established 3D convolutional neural network (CNN) architectures, including U-Net [10], VGG16 [2], and the simple VoxNet [34] algorithm, in order to benchmark our proposed method. Because these architectures operate on volumetric data, the original triangular meshes must be converted into 3D voxel grids prior to training. This section describes the preprocessing pipeline used to transform the electron microscopy (EM) triangular meshes into volumetric representations suitable for 3D CNNs.

S9.1. **Preprocessing Triangular EM Meshes into Volumes.** To apply 3D CNN architectures for segmentation, the triangular meshes must first be transformed into structured 3D voxel grids. This requires two main steps: resizing the mesh vertices and voxelizing the mesh into a multi-class 3D grid.

First, we resize each triangular mesh so that the resulting voxel grids remain computationally manageable. Larger meshes lead to significantly higher memory usage and computational cost; therefore, resizing the meshes according to hardware constraints is essential. In our experiments, each mesh is resized to around 5000 vertices. We also explored augmenting the dataset to increase variability, but the associated computational overhead made training prohibitively slow, and we therefore discontinued this approach.

Next, we convert the mesh into a volumetric representation using the `trimesh` voxelization pipeline. The voxelization step discretizes the mesh into a 3D grid, after which we fill the interior of the mesh using the `fill(method="orthogonal")` function. Voxels inside the dendritic branch are assigned label 1, while voxels outside remain 0.

After generating the dendritic branch volume, we construct the ground-truth shaft labels. To do this, we first create an expanded triangular mesh that tightly encloses the true shaft surface, by wrapping procedure based on Poisson surface procedure described in S6.1. We then voxelize and fill this expanded shaft mesh in the same manner as before. Voxels inside this expanded shaft region are assigned label 2. As the expanded mesh fully encloses the true shaft, it ensures that all shaft voxels are correctly captured, even in regions where the original mesh may be thin or irregular.

Finally, the shaft volume (label 2) is overlaid onto the dendritic branch volume (label 1), replacing any overlapping voxels. The resulting multi-class volume contains:

- label 0: background,
- label 1: dendritic spines,
- label 2: dendritic shaft.

Figure S6 depicts the voxel grid of a training dendrite, where three slices of the volume show the voxel grids in the x, y, and z planes. The background is labeled in white, the spines in blue, and the expanded shaft region in red.
This volumetric representation serves as the input to the 3D CNN architectures during training. The learning objective is to correctly identify the shaft voxels (label 2) within the dendritic branch volume. After training, the predicted shaft voxels can be mapped back onto the original triangular mesh to recover the segmented shaft surface.

S9.2. **Training Procedure.** The training procedure for the 3D CNN follows the same general strategy used for the DNN, with the main differences arising from the 3D convolutional architecture. We follow the same procedure described in Section S2. The strong class imbalance between shaft and spine vertices was handled using class weights computed from the inverse log-frequency of each class, which reduces bias toward the majority class and stabilizes training.
The network produces a volumetric prediction in which each voxel contains a vector of probabilities corresponding to the shaft and spine classes. A softmax activation function is applied to obtain voxel-wise class probabilities.
Because the final segmentation is defined on the mesh rather than the voxel grid, each mesh vertex is mapped to its corresponding voxel index. We then retrieve the probability values at that voxel location and use them to assign a class label to the mesh vertex, ensuring consistency between the volumetric CNN output and the surface-based representation of the dendrite. Since the meshes are downsampled before being fed to the CNN, the predicted probabilities are subsequently remapped back to the original high-resolution mesh. This yields the probability matrix $\overline{\mathbf{Y}} = f_\theta(\mathbf{Z}) \in [0,1]^{n\times 2}$, after which spine and shaft detection is restarted following the procedure described in Section 2.4.

### Supplemental Tables

| **Symbol** | **Meaning** |
|---|---|
| $\mathbf{X}_l$ | Vertex position on dendritic mesh |
| $\mathbf{E}_{l,m}$ | Edge vector between vertices $\mathbf{X}_l$ and $\mathbf{X}_m$ |
| $\mathbf{n}_j$ | Triangle normal vector |
| $A$ | Triangle area |
| $\mathbf{H}$, $\mathbf{K}$ | Mean and Gaussian curvature (smoothed) |
| $\widetilde{\mathbf{H}}$, $\widetilde{\mathbf{K}}$ | Sigmoid-enhanced curvature values |
| $a_{\mathbf{H}}, b_{\mathbf{H}}$; $a_{\mathbf{K}}, b_{\mathbf{K}}$ | Curvature-enhancement parameters |
| $k_{\text{bend}}$ | Bending coefficient in smoothing flow |
| $\mathbf{W}_{bend}$, $\mathbf{F}_{bend}$ | Bending energy and force |
| $\zeta(x)$ | Sigmoid function |
| $\mathbf{V}_j$ | Shaft-skeleton vertex |
| $\mathbf{D}_l$, $\mathbf{D}$ | Distance to nearest skeleton point; distance vector |
| $\mathbf{S}^k$ | K-means region label for $k$ clusters |
| $\mathbf{DNN}_1$, $\mathbf{DNN}_2$, $\mathbf{DNN}_3$ | Deep neural networks for segmentation |
| $\mathbf{Z}_{i,j}$ | Feature vector for vertex $i$ of mesh $j$ |
| $\mathbf{Y}_{i,j}$ | Ground-truth label (shaft/spine) |
| $f_\theta$ | Neural network mapping features to predictions |
| $\overline{\mathbf{Y}}$ | Predicted probability matrix |
| $\widetilde{\mathcal{X}}_{\text{spine}}$, $\widetilde{\mathcal{X}}_{\text{shaft}}$ | Vertices classified as spine / shaft |
| $a, b$ | Classification threshold parameters |
| $\mathcal{N}_i$ | 1-ring neighborhood of vertex $\mathbf{X}_i$ |
| $\mathcal{G}_i$ | Connected vertex group containing $\mathbf{X}_i$ |
| $\widetilde{\mathcal{Y}}^i_{\text{part}}$ | Connected component of a part type |
| $\widetilde{\mathcal{Y}}_{\text{shaft}}$ | All connected shaft components |
| $\mathcal{Y}^0_{\text{shaft}}$ | Largest shaft component (entire shaft) |
| $\mathcal{Y}_{\text{spine}}$ | Set of all spine components |
| $\mathcal{Y}^i_{\text{spine}}$ | Individual spine component |

Table S1. Summary of symbols and variables used in Sections 2.

| **Model** | **Runtime (s/Iterations)** | **Peak Memory (GB)** |
|---|---|---|
| $DNN_1$ | 47–50 | 0.114 |
| $DNN_2$ | 56–58 | 0.143 |
| $DNN_3$ | 43–45 | 0.314 |
| U-Net | 330–350 | 0.175 |
| VoxNet | 290–295 | 0.189 |
| VGG16–FCN | 370–380 | 0.211 |

Table S2. Runtime and memory benchmarks for all models. All training was performed on an HPCC CPU-only node with 2×Intel Cascade Lake CPUs (40 cores) and 754 GB RAM. All testing was performed on a MacBook Pro with an Apple M4 Max chip and 64 GB memory.

| **Category** | **Description** |
|---|---|
| **Training dataset** | Six high-resolution 3D EM reconstructions of dendritic segments from CA1 hippocampus of P21 rats. Slices received either control stimulation or *in vitro* theta-burst stimulation to induce LTP [7–9]. |
| **Training annotations** | Spines and shafts provided as separate watertight `.obj` meshes. Wrapped into unified dendritic meshes using the method in S6.1. |
| **Testing dataset** | Nanoconnectomic 3D EM reconstruction of hippocampal neuropil from the axon–spine coupling study [3]. |
| **Testing annotations** | 151 meshes available; 28 spiny dendritic branches selected based on annotation consensus. A mesh was included only if at least two annotators agreed on the presence of a spine. Individual spines retained only if labeled by at least two annotations. |
| **Visualization dataset** | Dendritic segments from the large-scale nanoconnectomic EM reconstruction of mouse neocortex by Kasthuri et al. [25]. This dataset provides densely reconstructed neuropil at nanometer resolution but does not include manual spine or shaft annotations. Used only for qualitative visualization (Figure 7). |
| **Visualization usage** | Used solely to demonstrate generalization of the method and to visualize predicted shafts and spines. Not used for training, validation, or quantitative testing. |

Table S3. Consolidated summary of datasets, annotations, and selection criteria used for training, testing, and qualitative visualization.

| Model | AUC | Criterion | Variant | Precision | Recall | F1-score |
|---|---|---|---|---|---|---|
| **U-Net** (a = 0.05) | 0.792 | IoU | Standard | 0.778 | 0.697 | 0.735 |
| | | | Union | 0.792 | 0.756 | 0.773 |
| | | DICE | Standard | 0.798 | 0.785 | 0.791 |
| | | | Union | 0.809 | 0.844 | 0.826 |
| **DNN$_3$** (a = 13) | 0.717 | IoU | Standard | 0.804 | 0.794 | 0.799 |
| | | | Union | 0.818 | 0.873 | 0.845 |
| | | DICE | Standard | 0.811 | 0.832 | 0.821 |
| | | | Union | 0.822 | 0.895 | 0.857 |
| **Vox Net** (a =10) | 0.682 | IoU | Standard | 0.817 | 0.311 | 0.451 |
| | | | Union | 0.811 | 0.299 | 0.437 |
| | | DICE | Standard | 0.840 | 0.366 | 0.510 |
| | | | Union | 0.836 | 0.355 | 0.498 |
| **DNN$_2$** (a = 0.5) | 0.605 | IoU | Standard | 0.805 | 0.667 | 0.729 |
| | | | Union | 0.824 | 0.754 | 0.788 |
| | | DICE | Standard | 0.813 | 0.702 | 0.753 |
| | | | Union | 0.833 | 0.804 | 0.818 |
| **VGG16 FCN** (a = 16) | 0.466 | IoU | Standard | 0.579 | 0.202 | 0.299 |
| | | | Union | 0.610 | 0.229 | 0.333 |
| | | DICE | Standard | 0.680 | 0.312 | 0.428 |
| | | | Union | 0.817 | 0.246 | 0.378 |
| **DNN$_1$** (a = 3) | 0.408 | IoU | Standard | 0.782 | 0.654 | 0.713 |
| | | | Union | 0.790 | 0.685 | 0.734 |
| | | DICE | Standard | 0.803 | 0.741 | 0.770 |
| | | | Union | 0.813 | 0.790 | 0.801 |

Table S4. Comparison of segmentation performance across geometric-informed models (**DNN$_1$**, **DNN$_2$**, **DNN$_3$**) and established CNN baselines (U-Net, VoxNet, VGG16-FCN). AUC is reported once per model, while Precision, Recall, and F1-score are evaluated under both IoU and DICE criteria, each with Standard and Union variants. Among the CNN baselines, U-Net achieves the highest AUC (0.792) and strong overall performance, whereas VoxNet and VGG16-FCN show moderate and weak performance, respectively. Across all metrics, the geometric-informed model **DNN$_3$** provides the most consistent and accurate segmentation, achieving the highest Recall and F1-scores—particularly under the Union variants, which better capture continuous spine regions. The weighting parameters used in the geometric-informed models are specified in (6), where the values of $a$ for each model are listed and the parameter $b$ is fixed at 0.81.

### Supplemental Figures

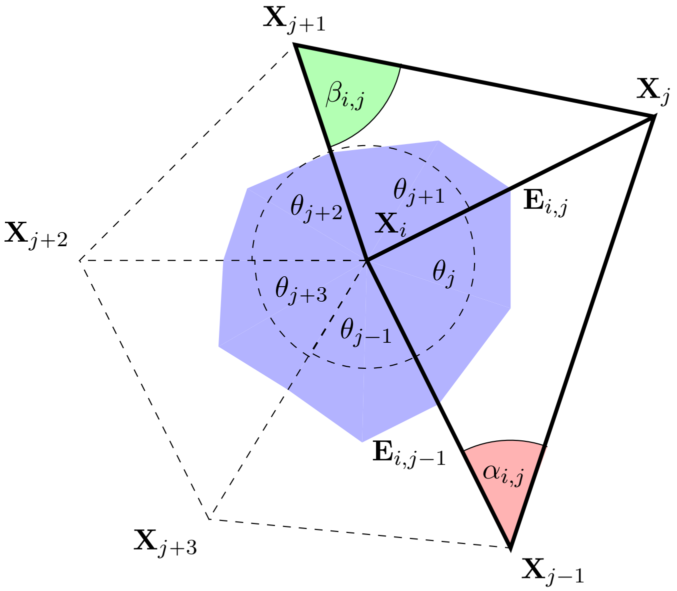


Figure S1. The graph shows the five triangular meshes of 1-ring neighbors to the point $p = \mathbf{X}_i$. It also depicts the incident angles $\alpha_{i,j}$ and $\beta_{i,j}$ opposite to the edge $\mathbf{E}_{i,j} = \mathbf{X}_j - \mathbf{X}_i$. The blue area constitutes the barycentric area $A_p$.

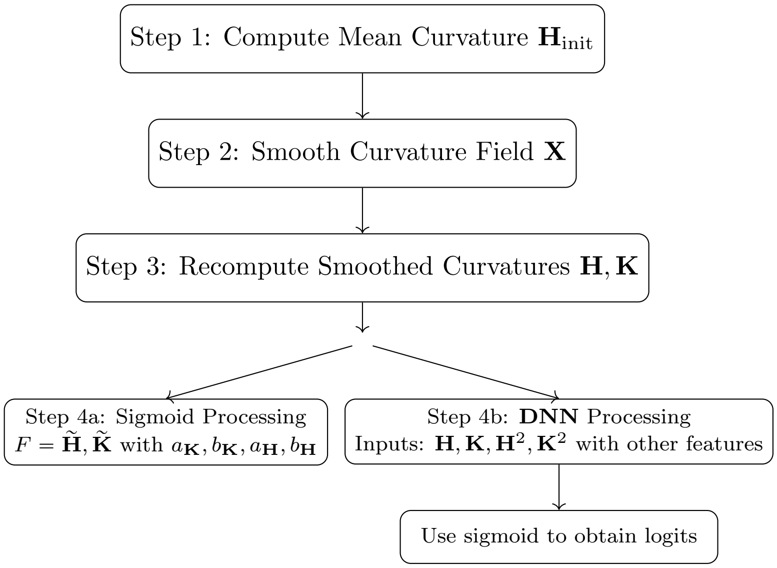


Figure S2. Flowchart of the mesh curvature preprocessing steps.

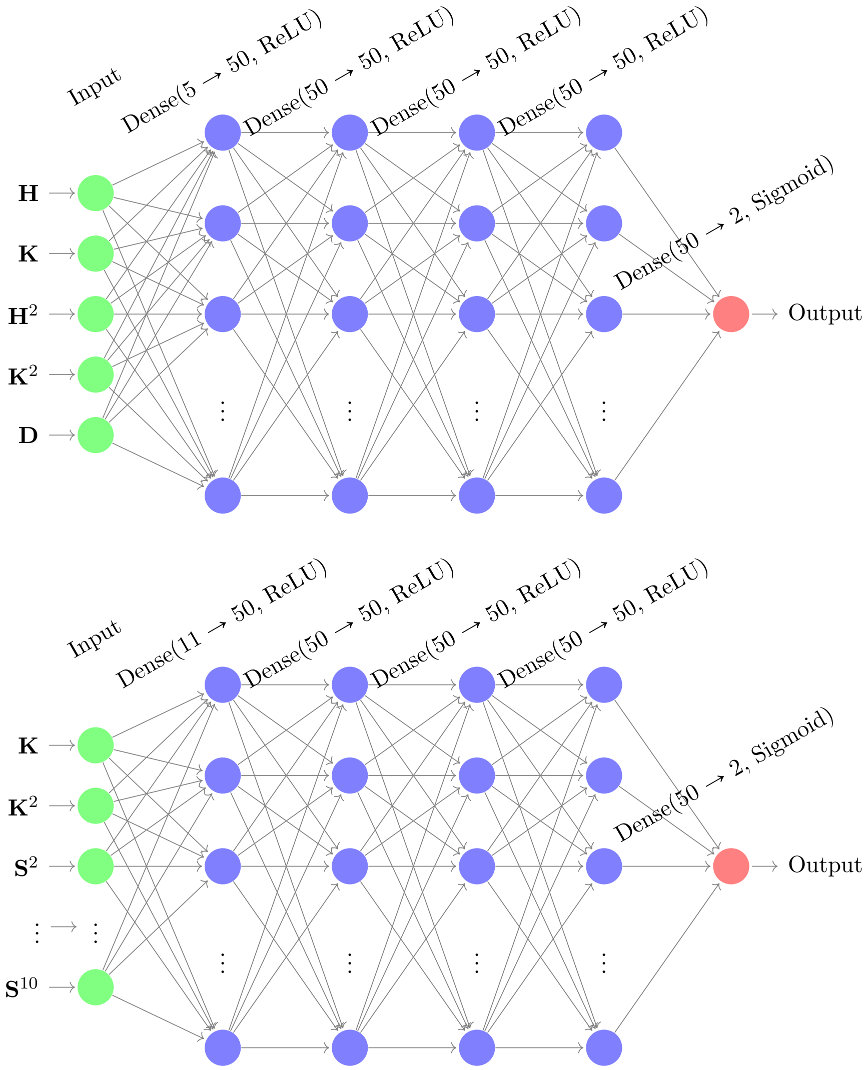


FIGURE S3. The top diagram illustrates the architecture of the deep neural network ($\mathbf{DNN}_2$), which improves upon $\mathbf{DNN}_1$. This model closely resembles the previous network but incorporates an additional input feature: the distance $\mathbf{D}$ between the central curve of the shaft (computed using $\mathbf{DNN}_1$) and the mesh vertices. The output layer employs a sigmoid activation function, consistent with the design of the earlier network. The bottom diagram shows the architecture of the deep neural network ($\mathbf{DNN}_3$) with additional input features. This model extends the previous design by enriching the input layer with multiple geometric and topological descriptors, including the Gaussian curvature and its squared value, as well as segmentation descriptors $\mathbf{S}^k$ obtained from K-means clustering of the shortest distances between mesh vertices and the dendrite skeleton. As in the earlier models, the output layer uses a sigmoid activation function.

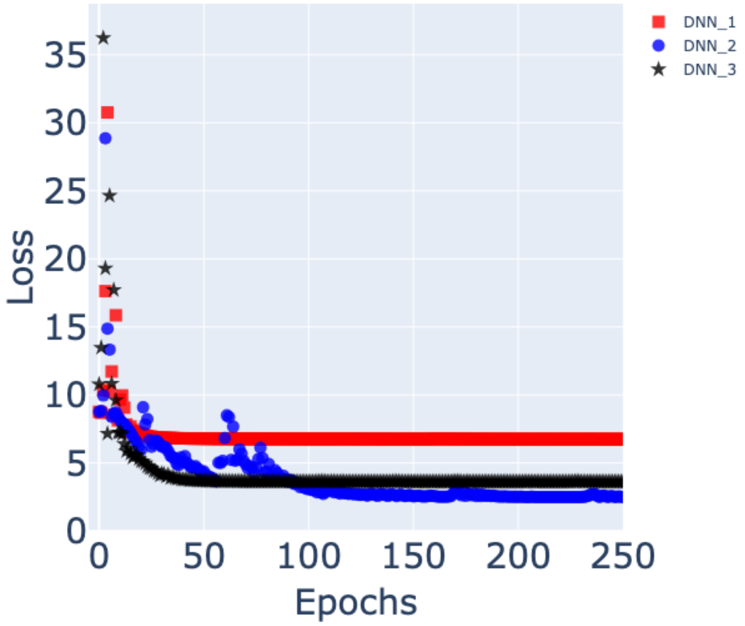


Figure S4. Training loss curves for the three deep neural network models. The red square markers, blue circular markers, and black star markers correspond to the loss curves of $\mathbf{DNN}_1$, $\mathbf{DNN}_2$, and $\mathbf{DNN}_3$, respectively. Among the three models, $\mathbf{DNN}_2$ achieves the best convergence, reaching a minimum loss of approximately 2.44, closely followed by $\mathbf{DNN}_3$ with a minimum loss of about 3.56. In contrast, the basic architecture $\mathbf{DNN}_1$ shows the poorest convergence behavior, with a substantially higher final loss of 6.73 throughout training. These results highlight the effectiveness of the additional features and architectural refinements introduced in $\mathbf{DNN}_2$ and $\mathbf{DNN}_3$ for improving model optimization and stability. The triangular mesh is obtained from CA1 dendritic reconstructions [7–9].

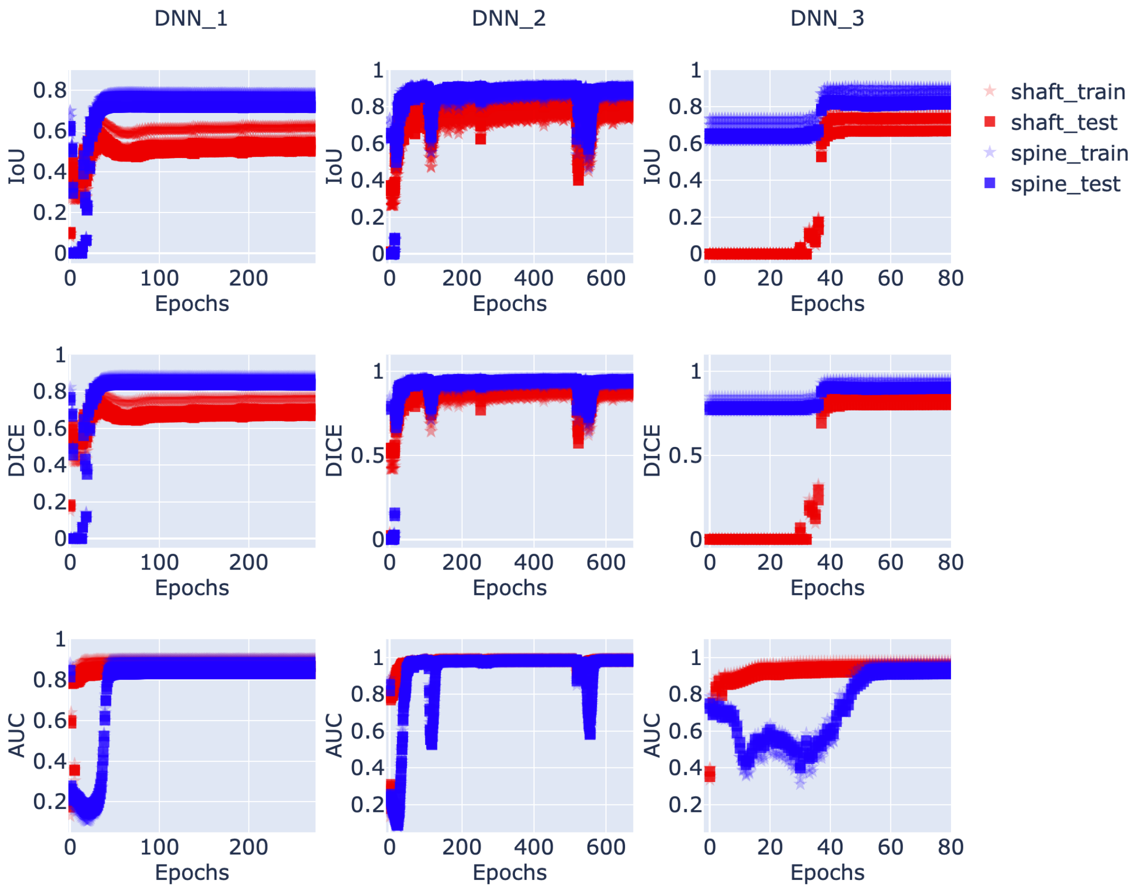


Figure S5. Performance curves for the three neural network models across the IoU, DICE, and AUC metrics. Columns 1, 2, and 3 correspond to $\mathbf{DNN}_1$, $\mathbf{DNN}_2$, and $\mathbf{DNN}_3$, respectively, and each row reports a different evaluation metric. Red curves represent dendritic shaft predictions and blue curves represent spine predictions; star markers denote values computed on the *training* datasets, while square markers indicate the corresponding *validation* datasets, all derived from [7–9]. All curves are computed exclusively from the *non-augmented* data. Overall, $\mathbf{DNN}_2$ and $\mathbf{DNN}_3$ exhibit very similar performance, with average IoU values of approximately 0.70 (shaft) and 0.81 (spines), and average DICE values around 0.88 (shaft) and 0.92 (spines). In contrast, $\mathbf{DNN}_1$ performs noticeably worse, particularly for shaft segmentation. AUC values remain high for all three models, with $\mathbf{DNN}_2$ and $\mathbf{DNN}_3$ again showing the strongest and most stable convergence. The close alignment between training and validation curves across all metrics indicates good generalization and minimal overfitting.

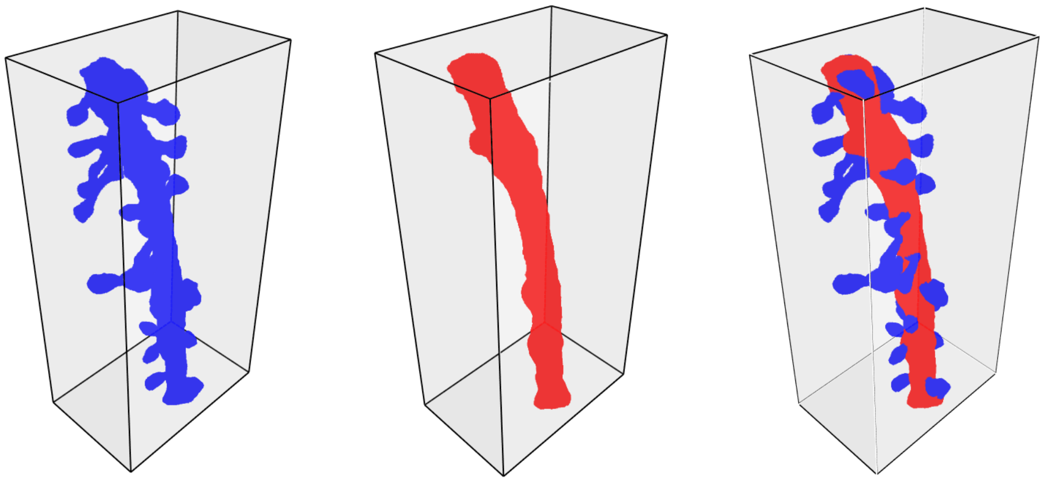

Figure S6. Construction of the voxel representations used for training. The top-left panel shows the voxelized triangular mesh of a dendritic branch (blue). The top-middle panel shows the ground-truth shaft labels generated by extending the annotated shaft mesh (red). The top-right panel shows the final target voxel grid, created by merging the dendrite-branch and shaft-label voxelizations and assigning the shaft label to overlapping voxels. This results in spine voxels labeled in blue and shaft voxels labeled in red. The bottom panel shows a slice view of the resulting volume in the plane defined by the two most informative PCA axes (PC1-PC2). The triangular mesh is obtained from CA1 dendritic reconstructions [7–9].
